# Clustered macrophages cooperate to eliminate tumors via coordinated intrudopodia

**DOI:** 10.1101/2024.09.19.613918

**Authors:** Lawrence J. Dooling, Alişya A. Anlaş, Michael P. Tobin, Nicholas M. Ontko, Tristan Marchena, Maximilian Wang, Jason C. Andrechak, Dennis E. Discher

## Abstract

Macrophages often pervade solid tumors, but their nearest neighbor organization is understudied and potentially enables key functions such as phagocytosis. Here, we observe dynamic macrophage clusters in tumors under conditions that maximize cancer cell phagocytosis and use reductionist approaches to uncover pathways to cluster formation and roles for tumor-intrusive pseudopodia, which we term ‘intrudopodia’. Macrophage clusters form over hours on low-adhesion substrates after M1 polarization with interferons, including T cell-derived cytokines, and yet clusters prove fluid on timescales of minutes. Clusters also sort from M2 macrophages that disperse on the same substrates. M1 macrophages upregulate specific cell-cell adhesion receptors but suppress actomyosin contractility, and while both pathways contribute to cluster formation, decreased cortical tension was predicted to unleash pseudopodia. Macrophage neighbors in tumor spheroids indeed extend intrudopodia between adjacent cancer cell junctions – at least when phagocytosis conditions are maximized, and coordinated intrudopodia help detach and individualize cancer cells for rapid engulfment. Macrophage clusters thereby provide a cooperative advantage for phagocytosis to overcome solid tumor cohesion.

## Introduction

Macrophages are tissue resident immune cells that often increase in number in pathologies such as cancer. Varied ontogenies and diverse stimuli contribute to phenotypic heterogeneity of macrophages in human cancers based on single-cell technologies (1–3). Although understanding is often limited, macrophage subtypes are distinguished based on surface marker expression, cytokine secretion profiles, and enzymatic activity. Differences in cell morphology (4, 5) and motility (6–8) have likewise been reported and point to underlying changes in adhesion receptors, cytoskeleton, and plasma membrane that also influence the defining function of a macrophage – phagocytosis (9–13).

Adhesion receptors and cytoskeleton are key to the dynamic interactions of a cell with its neighboring cells and with extracellular matrix. In solid tissues, macrophages make direct contact with numerous cell types including other macrophages. Clusters or aggregates of macrophages including fused multinucleated giant cells are often observed in response to large, nondegradable foreign bodies such as helminths and biomaterial implants as well as in granulomas during certain infections (14). In addition to these classic phenomena, macrophage aggregates and syncytia continue to be identified in new contexts including wound healing and fibrosis, neurodegeneration, and cancer (1, 15–18). In human head and neck cancer, macrophage subtypes identified by single-cell RNA sequencing as having either high *CXCL9* expression or high *SPP1* expression were described as accumulating as ‘clusters’ around tumor nests (1). Although cell-cell interactions and cell densities were unstudied, the anti-tumor *CXCL9*^+^ macrophages correlated with T cell infiltration, interferon-γ (IFNγ) signaling, and favorable patient outcomes whereas the pro-tumor *SPP1*^+^ macrophages associated with hypoxia and poor patient survival. Perivascular ‘nests’ of macrophages in murine breast tumors that depended on interleukin-6 and CCR5 were similarly associated with poor responses to chemotherapy and exclusion of CD8^+^ T cells (17). Earlier studies in human gastric cancer and endometrial cancer also reported improved survival of patients with ‘aggregated’ macrophages in tumor nests (19, 20). However, these qualitative descriptions of macrophage spatial organization do not explain how clusters or aggregates form in tumors or how macrophage clustering contributes to anti-tumor activity.

Many cell types retain the capacity for multicellular organization in vitro, which is perhaps best illustrated by sorting of cell mixtures in co-culture aggregates (21–24). Differential adhesion and differential contractility among the different cell types provide cogent explanations for demixing and sorting (25). However, macrophages and other immune cells have not been studied in such systems despite the importance of macrophage multicellular aggregates in physiological and pathological processes. Here, we investigated macrophage organization in response to various immune stimuli including polarizing cytokines and activation of Fcγ receptors during phagocytosis (26, 27). We show that different macrophage subtypes will cluster to varying extents in ‘immuno-tumoroid’ mixtures with cancer cells, as well as in monocultures. We discover and evaluate differential expression of key adhesion receptors among macrophages and also a counter- intuitive role for the actomyosin cytoskeleton and nucleoskeleton system in macrophage cluster formation. Live imaging of phagocytosis in tumor spheroids shows that neighboring macrophages extend intrusive pseudopods or ‘intrudopodia’ that coordinate with each other to disrupt tumor cohesion and thereby target individualized cancer cells for engulfment.

## Results

### Macrophages cluster under pro-phagocytic conditions in tumors and tumoroids

To investigate the spatial organization of macrophages in a therapeutically relevant tumor model, we generated lung metastases of syngeneic B16 melanoma cells and treated tumor-bearing mice with intravenous injections of an opsonizing IgG antibody targeting the melanocyte-specific membrane antigen Tyrp1 (**Fig.1a**). The B16 cells were engineered to lack CD47, a key inhibitory ligand that forms an immune checkpoint with signal regulatory protein α (SIRPα) on macrophages (28). We recently showed that anti-Tyrp1 injections reduce the number and size of B16 lung nodules and that responses are enhanced for CD47 KO B16 compared to wild-type (WT) B16 (29). Following the completion of the antibody treatment regimen, we harvested the lungs of treated mice and untreated control mice and stained for the mouse macrophage marker F4/80 (Fig.1a, **Fig.1a,b**). We found that macrophages in tumor nodules from treated mice were most often present in clusters, which we quantified as containing *N* = 1, 2, 3, 4… macrophages, whereas macrophages in tumor nodules from untreated control mice were most frequently present as isolated cells (i.e. *N* = 1) (**Fig.1b**, **Fig.1c**). From the distribution of cluster sizes *N* (Fig.1c), we computed the mean cluster size (*N*avg) and the *N*-weighted mean cluster size (*W*avg) for five nodules each from treated and untreated mice. Both methods of averaging revealed a statistically significant difference in the mean cluster size (Fig.1b). There was also a statistically significant difference in the cumulative distributions of *N* across all treated nodules versus untreated nodules (Fig.1c, inset). Importantly, macrophages in clusters frequently appeared to be phagocytosing B16’s based on the internalization of both large nuclei characteristic of B16’s and melanin pigments visible in bright-field (Fig.1a). Together, these results suggest an association between macrophage clustering and phagocytosis in tumors.

**Figure 1.**
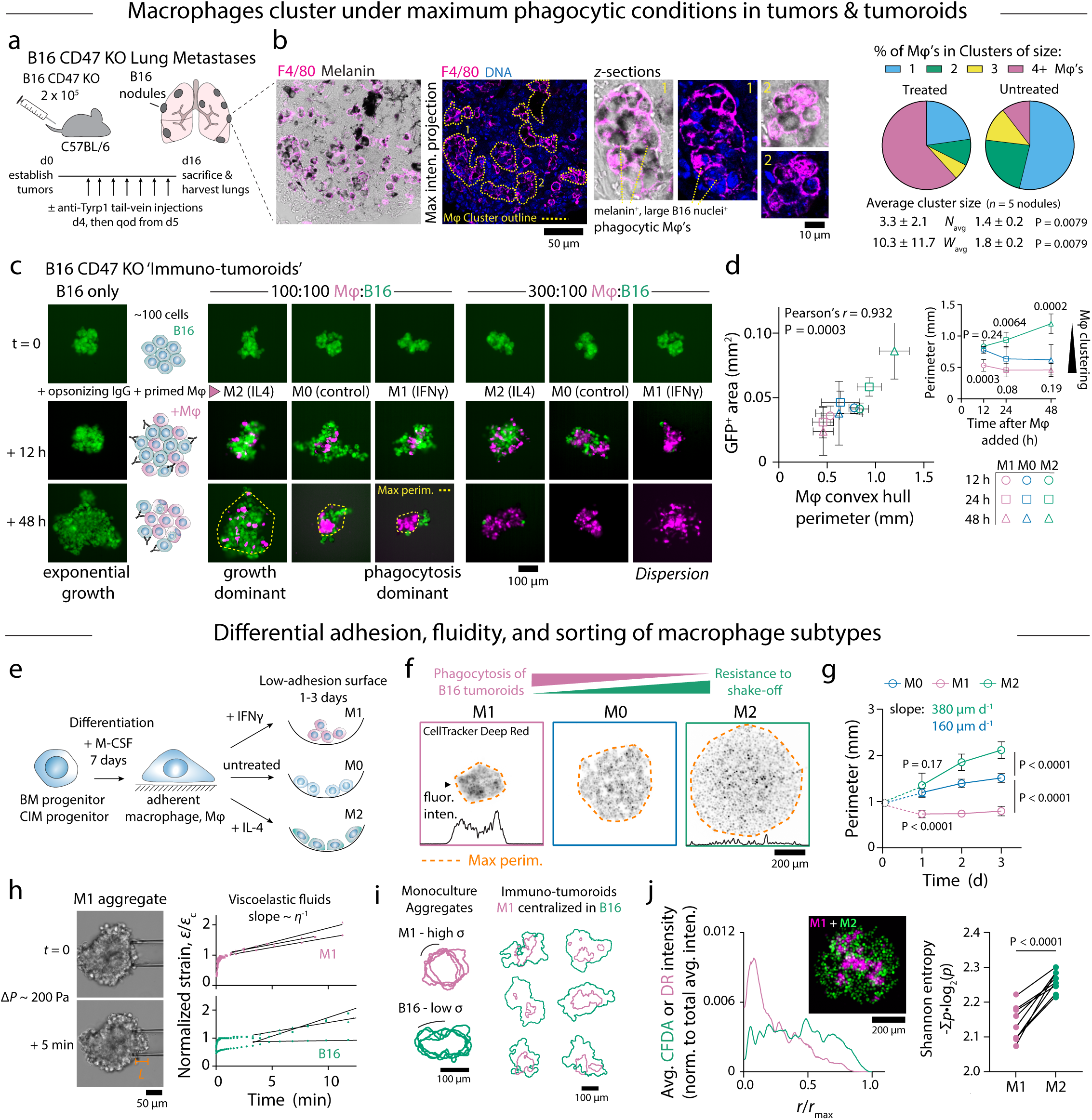
Macrophages cluster under maximum phagocytic conditions and when polarized with interferon-γ (IFNγ). a. Establishing and treating B16 CD47 KO lung metastases in mice. b. Bright-field and confocal fluorescence microscopy of a B16 CD47 KO metastatic nodule immunostained for F4/80. The yellow dotted outlines denote macrophage clusters. Scale bar 50 μm. Zoomed confocal slices of two macrophage clusters. Scale bar 10 μm. Pie-charts: Percentage of macrophages that are part of clusters of *N* = 1, 2, 3, and 4 or more macrophages in nodules from treated and untreated mice. Number average cluster size (*N*avg) and *N*-weighted average cluster size (*W*avg) in nodules from treated and untreated mice. Mean ± standard deviation (s.d.), *n* = 5 nodules per condition, Mann-Whitney test (two-tailed). c. Immuno-tumoroids comprising growing aggregates of B16 CD47 KO cells (green) to which macrophages (Mφ’s, magenta) primed with IFNγ or interleukin-4 (IL4) were added at varying ratios with an opsonizing antibody (anti-Tyrp1). The yellow dotted outlines depict the perimeter of a 2D convex hull enclosing all macrophages. Scale bar 100 μm. d. The projected GFP^+^ area of B16 CD47 KO in immuno-tumoroids plotted against the macrophage convex hull perimeter for different time points and cytokine priming conditions. Inset: Convex hull perimeter as a function of time after macrophages were added. Mean ± s.d., *n* = 7 or 8 wells per condition, repeated measures two-way ANOVA with Dunnett’s multiple comparisons test. e. Macrophage differentiation from bone marrow (BM) or conditionally immortalized macrophage (CIM) progenitors and clustering assay on low-adhesion U-bottom well plates. f,g. Representative fluorescence images after 3 days (**f**) and convex hull perimeters as of function of time (**g**) of macrophages on low-adhesion surfaces in the presence of 20 ng mL^-1^ IFNγ (= M1), no cytokine (= M0), or 20 ng mL^-1^ IL4 (= M2). The orange dotted outlines depict the 2D convex hull enclosing all macrophages. The fluorescence intensity profile corresponds to a line across the image at the position denoted by the arrowhead. Scale bar 200 μm. Mean ± s.d., *n* = 9 or 10 wells per condition, repeated measures two-way ANOVA with Dunnett’s multiple comparisons test. h. Bright-field images of micropipette aspiration of an M1 macrophage cluster and creep profiles of M1 macrophage cluster (*top*) and B16 tumoroid (*bottom*) aspiration. The strain (*ε* = aspiration length *L* / pipette radius *R*P) is normalized by a critical strain *ε*c corresponding to the transition to the viscous regime. Solid lines are linear fits of the viscous regime where the slope is inversely proportional to viscosity, *η*. Scale bar 50 μm. i. Outlines of M1 macrophage clusters and B16 tumoroid clusters (4 each) and outlines of B16 and M1 macrophages in immuno-tumoroids at *t* = 12 h depicting the centralization of macrophage clusters within B16 tumoroids. Scale bar 100 μm. j. Radial fluorescence intensity profiles and fluorescence image of a representative co-culture comprising M1 (magenta, CellTracker Deep Red) and M2 (green, CFDA) macrophages in a low-adhesion U-bottom well after 1 day. Scale bar 200 μm. Shannon entropy computed from the M1 and M2 radial profiles for *n* = 8 different co-cultures, Student’s *t*-test (two-tailed, paired).

Recently described ‘clusters’ and ‘nests’ of macrophages in human and murine tumors have been related to pro-inflammatory signaling pathways activated in subsets of tumor macrophages (1, 17). To begin to address the effects of tumor macrophage heterogeneity on clustering and phagocytosis, we performed multi-day ‘immuno-tumoroid’ assays in which mouse bone marrow-derived macrophages (BMDMs) were activated with IFNγ (‘M1’) or interleukin-4 (IL4, ‘M2’) and added together with anti-Tyrp1 opsonizing IgG to cohesive aggregates of B16 cells on low-adhesion, U-bottom well plates (30). We confirmed BMDM expression of macrophage markers including FcγRI and SIRPα, which positively and negatively regulate antibody-dependent cellular phagocytosis, respectively (**Fig.2a**). We also confirmed the expected upregulation of MHCII and CD206 on BMDMs that were ‘primed’ for two days in standard 2D cell culture conditions with IFNγ and IL4, respectively (**Fig.2b,c**). We then added primed macrophages to preformed B16 CD47 KO tumoroids and quantified the GFP^+^ area versus time as a proxy for the number of B16 cells (**Fig.1c**). The resulting growth curves were fit to a simple exponential model with an effective growth rate, *k*eff, that is positive for net B16 growth and negative for net repression by macrophages. Consistent with our earlier study (30), elimination of B16’s by unprimed control (M0) macrophages required opsonization with anti-Tyrp1, and the nonlinear dependence of *k*eff on the number of added macrophages was well fit by a cooperative phagocytosis model (**Fig.3a**). Tumoroid growth suppression was enhanced by M1-priming: although still cooperative with respect to the number of macrophages added, *k*eff was lower (i.e. faster elimination) with M1- primed macrophages than an equivalent number of unprimed macrophages and there was a shorter delay time (Fig.3a). Additionally, the requirement for anti-Tyrp1 was relaxed, and M1- primed macrophages could eliminate unopsonized tumoroids when added at a 3:1 or 9:1 ratio to B16’s whereas M0 macrophages could not (Fig.3a). Elimination of B16 tumoroids with WT CD47 levels still required opsonization with anti-Tyrp1 regardless of macrophage priming, implicating phagocytosis that is inhibited by CD47-SIRPα as the predominant effector function (**Fig.3b**). M2-primed macrophages were less effective against tumoroids than either M1- or M0-primed macrophages (Fig.1c, Fig.3a).

In addition to the differences in their ability to suppress tumoroid growth, M1- and M2-primed macrophages exhibited differences in clustering in immuno-tumoroids. This was most evident at a nominal BMDM:B16 ratio of 1:1 where M1-primed macrophages clustered even more strongly than M0 macrophages while M2 macrophages dispersed throughout the tumoroid. To quantify the extent of clustering or dispersion, we computed a convex hull – i.e. the smallest convex shape that encapsulates all macrophages – and used the convex hull perimeter as a metric for comparing different experimental conditions (Fig.1c yellow outline). The convex hull perimeter was smallest for M1-primed macrophages and largest for M2-primed macrophages (**Fig.1d**), and – importantly – M0 were statistically similar to M1 by this measure at 24 h and 48 h. The convex hulls of the M2-primed macrophages increased with time as the tumoroid expanded due to B16 proliferation that was not effectively suppressed by phagocytosis (Fig.1d inset). On the other hand, the convex hulls of M1-primed and M0 macrophages decreased in size over time, which likely reflects a combination of macrophage clustering and tumoroid shrinkage as phagocytosis dominates proliferation. From these observations, we conclude that macrophage priming with IFNγ enhances both macrophage clustering and elimination of B16’s from tumoroids while priming with IL4 has the opposite effect on both processes.

### Differential adhesion, fluidity, and sorting of macrophage subtypes

We next examined clustering and dispersion of M1 and M2 monocultures on the same low-adhesion well plates used for immuno-tumoroids as a further reductionist approach to understanding differences in macrophage spatial organization (**Fig.1e**). We observed that M1 macrophages form compact clusters whereas M2 macrophages instead disperse (**Fig.1f**), which is similar to their respective behaviors in immuno-tumoroids (Fig.1c). Unpolarized M0 macrophages have an intermediate phenotype and are more dispersed than M1 macrophages despite local regions where cells adhere to one another. To quantify these differences, we again computed the perimeter of a convex hull enclosing all cells (Fig.1f orange outline). The trend in the average perimeter – M1 < M0 < M2 – is roughly the same as for the immuno-tumoroids, except that M0 was much closer to M1 under conditions of phagocytosis (Fig.1d) – as emphasized. This trend is certainly consistent with the polarizing effects of IFNγ and IL4 that give rise to diametrically opposed macrophage states, which manifest here as a tendency to cluster or disperse, respectively. M1 clusters could be easily dislodged by inverting the well plate or by pipetting with a wide-bore tip but remained intact and cohesive during these manipulations. M2 macrophages tended to be more strongly attached to the substrate, although this depended somewhat on the specific surface treatment. M2 macrophages on surfaces passivated by poly(2-hydroxyethylmethacrylate) (pHEMA) tended to remain round while M2 macrophages on surfaces passivated by a commercial anti-adherence surfactant treatment were frequently observed to elongate with prominent lamellipodia, particularly at the leading edge (**Fig.4a**).

Macrophage polarization alters the expression of various chemokines and chemokine receptors, and it is conceivable that increased expression of chemokine-receptor pairs could create a scenario in which macrophages are attracted to one another leading to cluster formation, as recently proposed for CCR5-dependent macrophage nests in murine breast cancer (17). To assess this possibility, we performed our clustering assay with M0 and M1 macrophages in the presence of pertussis toxin, a potent inhibitor of G-protein receptor signaling and leukocyte chemotaxis (31). However, pertussis toxin had no effect on the ability of M1 macrophages to cluster (**Fig.4b**). Instead, treatment limited the dispersion of M0 macrophages resulting in a smaller convex hull perimeter. This suggests that chemokine signaling pathways are not required in M1 macrophage clustering but instead might contribute to the dispersion of M0 and M2 macrophages, perhaps through cell migration in response to gradients of macrophage colony-stimulating factor (M-CSF) or serum proteins generated by localized depletion in the center of the U-bottom well. This scenario would also be consistent with the migratory phenotype of macrophages at the leading edge of some dispersing M2 cultures (Fig.4a).

Kinetic studies revealed that M1 clusters form in less than 24 h while the dispersion of M2 and M0 macrophages occurs over the course of days (**Fig.1g**). For insight into clustering, we performed kinetic measurements of M0 and M1 macrophages over the first day following transfer of cells suspensions to low-adhesive substrates (**Fig.4c**). The perimeter of the convex hull enclosing cells decreased dramatically between 1 and 3 h under both M0 and M1 conditions, which likely reflects cell sedimentation or rolling along the substrate toward the center of the U-bottom well. A statistically significant difference between M0 and M1 macrophages was evident by 6 h after plating, and the two macrophage subsets diverged further by 24 h. The compaction of M1 macrophages into tight clusters requires de novo protein synthesis as it did not occur in the presence of the protein translation inhibitor cycloheximide (Fig.4c). Together, these results suggest that rapid changes in M1 macrophages including expression of new proteins drives them to adhere to one another in clusters.

To assess the capacity for repolarization, macrophages were primed in standard 2D culture for 2 days under M0, M1, or M2 conditions, detached to generate single-cell suspensions that were divided into three samples containing IFNγ, IL4, or no cytokine, and finally added to low-adhesion wells for a total of nine conditions (3 priming conditions x 3 assay conditions) (**Fig.4d**). Regardless of the priming condition, macrophages were always well compacted within 24 h in the presence of IFNγ and remained so for the duration of the experiment. On the other hand, M1- primed macrophages that were switched to M0 or M2 conditions were initially clustered at day 1 but gradually dispersed. The effects of M2 polarization leading to dispersion are slower than the aggregating effects of M1 polarization. There was no significant difference between M0 and M2 macrophages after 1 day on the low-adhesion substrates (Fig.1g), but M2 macrophages become more dispersed than M0 macrophages by the second day. Consistently, macrophages primed for 2 days with IL4 were significantly more dispersed than unprimed macrophages within 24 h of transfer to the low-adhesion substrate (**Fig.4e**), indicating they had enough time during the priming phase to upregulate pathways leading to dispersion. Among different mouse donors, we observed that M1-primed macrophages switched to M0 conditions were initially more compact when macrophages came from female mice than male mice, suggesting sex differences to the inflammatory memory of the priming period (**Fig.4f**). Overall, we conclude that the effects of polarizing cytokines on macrophage clustering and dispersion in our low-adhesion assay are reversible, but kinetics vary for transitions between different states and might also depend on donor sex.

Nearly identical clustering and dispersion behavior was observed using mouse macrophages differentiated from HoxB8-estrogen receptor conditionally immortalized macrophage progenitor cells (hereafter referred to as CIMs) (30, 32, 33). Differentiated CIMs expressed similar levels of myeloid and macrophage surface markers as BMDMs (Fig.2a) and could also be polarized to M1 and M2 states, although the basal and IFNγ-induced MHCII levels in differentiated CIMs were lower than in BMDMs (Fig.2b,c). Nonetheless, differentiated and polarized CIMs cultured on low-adhesion substrates showed the same trends as BMDMs, and the convex hull perimeters of the two types of macrophages under M0, M1, and M2 conditions were highly correlated with one another (**Fig.4g-i**).

To characterize the physical state of M1 clusters, we performed micropipette aspiration using pipette tips with diameters of approximately 50 μm, which is wide enough to accommodate many cells. M1 clusters behaved as viscoelastic liquids with a short-time elastic response that gave way to cohesive flow on longer time scales (**Fig.1h**), which is qualitatively similar to the behavior of B16 aggregates (30). M1 clusters generally have a smoother surface than the more fractal perimeters of B16 aggregates, suggesting a higher surface tension (*σ*). Consistent with previous descriptions of liquid-like tissue aggregates (22, 25), the higher *σ* macrophages are located more centrally than the lower *σ* B16’s in ‘immuno-tumoroid’ co-cultures (**Fig.1**c,**i**). Moreover, in mixtures of M1 and M2 macrophages labeled with different CellTracker fluorescent dyes, both states retained the behavior observed in monocultures, thereby producing a sorting effect in which M1 macrophages occupied the center of the co-culture and M2 macrophages predominated at the periphery (**Fig.1i**). From radial fluorescence profiles approximating the probability distributions of locating M1 and M2 macrophages in the mixture, we computed a Shannon entropy that was always lower for M1’s than M2’s (**Fig.1j**), indicating more order or structure in the spatial distribution of M1 macrophages.

The clustering of macrophages activated by a pro-inflammatory stimulus is at least somewhat generalizable. A second M1 stimulus, the toll-like receptor 4 agonist lipopolysaccharide (LPS), produced an effect similar to IFNγ on both BMDMs and CIMs, and there was no additive or synergistic effect of combining IFNγ and LPS (**Fig.2a**, **Fig.4j-l**). Interferon-α (IFNα) drove macrophage clustering but to a lesser extent than IFNγ or LPS. Macrophage clustering driven by IFNγ and IFNα were dose-dependent with EC50 values of ∼50 pg/mL and 1 pg/mL, respectively (**Fig.5a,b**), suggesting that specific signaling events downstream of receptor activation underlie the macrophage clustering observed here.

**Figure 2.**
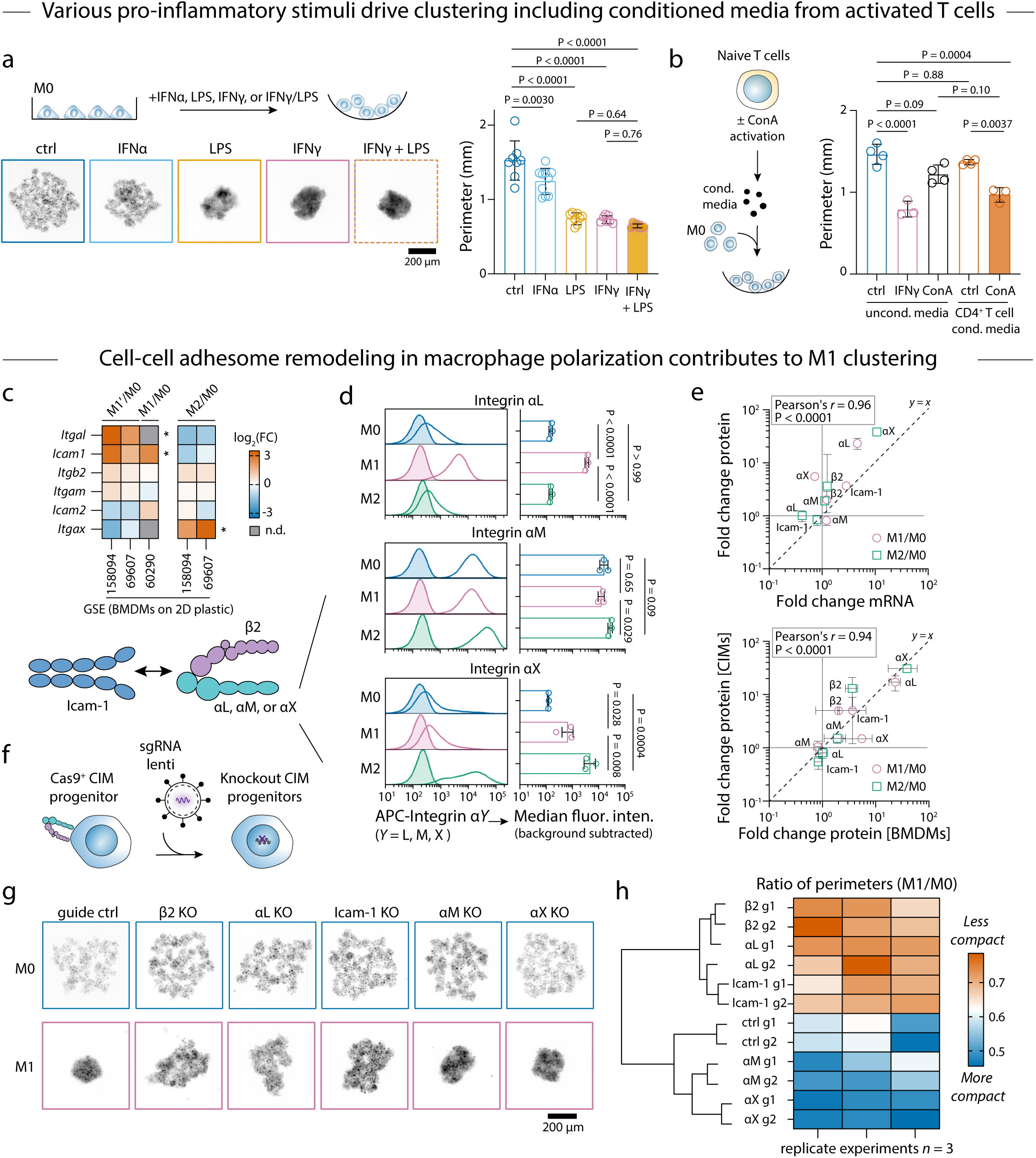
Macrophage clustering with pro-inflammatory stimuli and roles for differential expression of adhesion receptors a. Representative fluorescence images and convex hull perimeters of BMDMs after 1 day on low-adhesion U-bottom well plates in the presence of 20 ng mL^-1^ interferon-α (IFNα), 100 ng mL^-1^ lipopolysaccharide (LPS), 20 ng mL^-1^ IFNγ, or 20 ng mL^-1^ IFNγ + 100 ng mL^-1^ LPS. Mean ± s.d., *n* = 7-10 wells per condition, ordinary one-way ANOVA with Tukey’s multiple comparisons test. Scale bar 200 μm. b. Convex hull perimeters of BMDMs after 1 day on low-adhesion U-bottom well plates in the presence of conditioned media from concanavalin-A (ConA)-activated or unactivated mouse CD4^+^ T cells, or in unconditioned media with ConA or IFNγ. Mean ± s.d., *n* = 3 or 4 wells per condition, ordinary one-way ANOVA with Šidák’s multiple comparisons test between indicated groups. c. Heat map of fold changes (FC) in expression of genes encoding adhesion receptors in polarized (M1, M1’ or M2) vs. unpolarized (M0) BMDMs from datasets listed in Table S1. * indicates an adjusted P < 0.05 across all datasets in which the gene was detected (*top*). n.d., not detected. Macrophage integrin heterodimers comprising an αL, αM, or αX subunit and the β2 subunit interacting with an Icam-1 homodimer (*bottom*). d. Flow cytometry of M0, M1, and M2 BMDMs stained for integrin αL, αM, and αX. Histograms show staining of a representative BMDM culture and bar graphs report the median fluorescence intensity (MFI) of stained cells corrected by subtracting the MFI of unstained cells. Mean ± s.d., *n* = 3 BMDM cultures from different mice, ordinary one-way ANOVA and Tukey’s multiple comparisons test on log-transformed MFI values. e. Correlation between fold changes in surface protein expression (from flow cytometry in panel **d**) and fold changes in gene expression (from GSE69607 dataset in panel **c**). Correlation between fold changes in surface protein expression on BMDMs (panel **d**) and CIMs (**Fig.6d**). f. Generation of knockout CIM progenitor lines using lentiviral transduction of single guide RNA (sgRNA) constructs to Cas9^+^ CIMs. g. Representative fluorescence images of macrophages differentiated from guide control and KO CIM progenitors after 24 h on low-adhesion U-bottom surfaces under M0 and M1 conditions. Scale bar 200 μm. h. Heat map and hierarchical clustering dendrogram for the ratio of mean convex hull perimeters of M1 and M0 cultures on low-adhesion, U-bottom surfaces. *n* = 3 replicate experiments.

Lymphocytes are the most likely source of IFNγ in vivo, and lymphocyte secreted factors have long been known to promote macrophage aggregation (34). Therefore, we isolated naïve CD4^+^ and CD8^+^ T cells from mouse spleens and activated them ex vivo with concanavalin A (ConA) (**Fig.2b**, **Fig.5c**). Adding conditioned media from activated T cells to M0 macrophages resulted in statistically significant clustering based on measurement of the convex hull perimeters, although not to the extent observed with recombinant IFNγ. Neither conditioned media from naïve T cells nor an equivalent concentration of ConA added to unconditioned media had significant effects on clustering.

### Differences in Adhesion Receptor Repertoires in Polarized Macrophages

Differences in adhesion receptors could help to explain the clustering and sorting behavior of different macrophage subsets. Therefore, we analyzed public transcriptomic data for differential expression of genes encoding such receptors in BMDMs polarized to M1 or M2 states (**Table S1**). Of the three M1 datasets analyzed, one included macrophages polarized with IFNγ only for 18 h and the two others included macrophages polarized with IFNγ + LPS for 24 h. Because IFNγ + LPS produced a similar level of macrophage clustering as IFNγ alone (Fig.2a, Fig.2j-l), we presumed that any key differences in gene expression underlying this effect would be reflected in both conditions. Nonetheless, we refer to macrophages activated with IFNγ + LPS as M1’ macrophages to avoid confusion. In both M2 datasets analyzed, BMDMs were polarized with IL4 for 24 h.

Although multiple types of cell-surface receptors have been implicated in macrophage adhesion or fusion, we chose to focus on differential expression of integrins considering they mediate both cell-cell and cell-ECM interactions (**Fig.2c**). Macrophages express multiple integrins including heterodimers comprising αL, αM, αX, or αD subunits and the leukocyte-specific β2 subunit. Expression of *Itgam* and *Itgb2* encoding the αM and β2 subunits, respectively, did not change significantly upon M1 or M2 polarization, and *Itgad* encoding the αD subunit was of very low abundance in the RNAseq data and not detected either microarray dataset. On the other hand, expression of *Itgal* encoding the integrin αL subunit was upregulated >5-fold in the two M1’ datasets in which it was detected while expression of *Itgax* encoding the integrin αX subunit was upregulated >4-fold in both M2 datasets. The αLβ2 integrin (also known as LFA-1) is expressed widely across immune cells and best known to mediate cell-cell adhesions in the context of leukocyte transendothelial migration (35) and the immune synapse (36). Expression of *Icam1* encoding the integrin ligand intercellular adhesion molecule-1 (Icam-1) was also upregulated >3-fold in all three M1 datasets, providing a candidate receptor-ligand pair that could mediate homotypic cell-cell adhesion between M1 macrophages (Fig.2e, bottom).

Gene expression measurements in the M1’ and M2 RNAseq dataset were made at multiple time- points (37), allowing us to compare the kinetics of gene expression changes to the kinetics of macrophage clustering (Fig.4c). *Itgal* and *Icam1* expression reached peak levels after only 4-6 h in M1’ macrophages but remained elevated after 24 h relative to M0 macrophages (**Fig.6a**). Such a rapid change in the levels of genes encoding adhesion receptors is consistent with our studies of macrophage clustering kinetics in which statistically significant differences between M1 and M0 macrophages emerge within 6 h after plating in U-bottom wells (Fig.4c). The same rapid induction of *Itgal* and *Icam1* occurred following M2 to M1 repolarization (**Fig.6b**), which is also consistent with our clustering results with repolarized BMDMs (Fig.4d). Therefore, increased cell-cell adhesion mediated by increased levels of αLβ2 integrin and Icam-1 could account for M1 clustering, at least based on changes in gene expression.

We next verified that the gene expression changes reported by others were reflected in changes to surface protein levels in our polarized BMDMs. Flow cytometry revealed upregulation of integrin αL and Icam-1 in M1 macrophages and upregulation of integrin αX in M2 macrophages (**Fig.2d**, **Fig.6c**). There was a high degree of correlation between changes at the transcript level and surface protein level and also between surface protein level changes in BMDMs and CIMs (**Fig.2e**, **Fig.6d**). BMDMs treated with LPS for 24 h upregulated integrin αL and Icam-1 to a similar extent as BMDMs treated with IFNγ, although the change in Icam-1 was not statistically significant (**Fig.6e**). BMDMs treated with IFNα had only a small increase in integrin αL surface expression and no significant change in Icam-1 (Fig.6e). Overall, these measurements are consistent with the similar clustering observed for macrophages activated with IFNγ, LPS, or IFNγ + LPS, which was greater than clustering of macrophages activated with IFNα.

We undertook two approaches to determine the functional significance of differential integrin expression on macrophage clustering. First, we assessed clustering and dispersion of M0, M1, and M2 BMDMs in the presence of blocking antibodies against the αL, αM, and β2 integrin subunits and Icam-1 as well as anti-integrin β1 and isotype control antibodies. None of these blocking antibodies had a significant effect on M1 clustering (**Fig.7a**). Instead, anti-integrin αM and β2 drove tight clustering of both M0 and M2 BMDMs. Similar results were obtained with anti-integrin αM and β2 F(ab’)2 fragments, indicating macrophage Fc receptors were not likely not involved (**Fig.7b**). While such clusters could result from disrupting cell-substrate adhesions involving αMβ2 integrin, the relatively high expression of αM compared to αL and αX led us to instead suspect that the antibodies were agglutinating macrophages.

The inability of any blocking antibodies to prevent M1 clustering and the likely agglutinating activity of some anti-integrin antibodies prompted us to seek an alternative approach. We therefore took advantage of Cas9 nuclease expression in CIMs (33) to knock out the αL, αM, αX, or β2 integrin subunits or Icam-1. We generated lentiviruses to express single guide RNA (sgRNA) sequences targeting the genes encoding those proteins and transduced Cas9^+^ CIM progenitors to generate stable knockout (KO) lines (**Fig.2f**, **Fig.6f**). Clustering in the presence IFNγ was unaffected in αM KO’s or αX KO’s but significantly abrogated in αL, β2, or Icam-1 KO’s (**Fig.2g**). To quantify differences in M1 clustering, we computed the ratio of the average M1 and average M0 convex hull perimeters for each KO line (**Fig.2h**). Given that none of the KO’s exhibited obvious differences in the M0 state, a higher ratio indicates less compact clustering induced by M1 polarization whereas a lower ratio indicates more compact clustering. We performed hierarchical clustering of the M1/M0 ratios for the different cell lines and found the longest distance to be between the cluster containing the αL KO, β2 KO and Icam-1 KO lines and the cluster containing the αM KO, αX KO, and non-targeting guide control lines. These results strongly support a role for αLβ2 integrin binding to Icam-1 in mediating cell-cell adhesions among macrophages in M1 clusters.

### Upregulation of adhesion receptors under phagocytic conditions in tumoroids

Macrophage clustering in IgG-opsonized immuno-tumoroids is enhanced by M1-priming, but clustering is still observed with unprimed macrophages (Fig.1c,d). In addition to well-known effects on actin polymerization, Fc receptor activation initiates signaling that leads to transcriptional changes on the order of hours (27), a relevant timescale for immuno-tumoroids. This raises the possibility that signaling associated with phagocytosis upregulates surface receptors to promote macrophage cell-cell adhesion as observed with IFNγ- and LPS-treated macrophages. To investigate this, we collected and pooled macrophages from immuno-tumoroids after 18 h and stained them for integrin αL and Icam-1 as well as MHCII. As expected from our recent study (30), there were significantly more phagocytic macrophages (CD45^+^GFP^+^) in opsonized immuno-tumoroids compared to unopsonized immuno-tumoroids (**Fig.3a,b**, **Fig.8a,b**). Parallel cultures in which macrophages were fluorescently labeled with a CellTracker dye and added to immuno-tumoroids for imaging over the course of several days revealed a near-complete elimination of B16’s from opsonized immuno-tumoroids with evidence for macrophage clustering followed by dispersion based on the convex hull perimeters (**Fig.8c,d**). However, the kinetics for elimination, clustering, and dispersion varied considerably from well to well, which is expected to introduce significant variability in measurements of pooled immuno-tumoroids. Nonetheless, we proceeded to compare integrin αL, Icam-1, and MHCII surface expression on phagocytic macrophages versus non-phagocytic macrophages (CD45^+^GFP^-^). The histogram distributions of fluorescence intensity levels were shifted rightward toward higher values for all three markers on phagocytic macrophages relative to non-phagocytic ones (**Fig.3c**). Considering these rightward shifts, we plotted the median, 75^th^ percentile, and 95^th^ percentile values against the percentage of phagocytic macrophages and determined that the 95^th^ percentile of Icam-1 and integrin αL and the 75^th^ percentile of Icam-1 are significantly correlated with the level of phagocytosis (**Fig.3d**). Together, these findings suggest that phagocytic conditions in immuno-tumoroids might enhance macrophage clustering in part through increased surface expression of the adhesion receptors αLβ2 and Icam-1 (**Fig.3e**).

**Figure 3.**
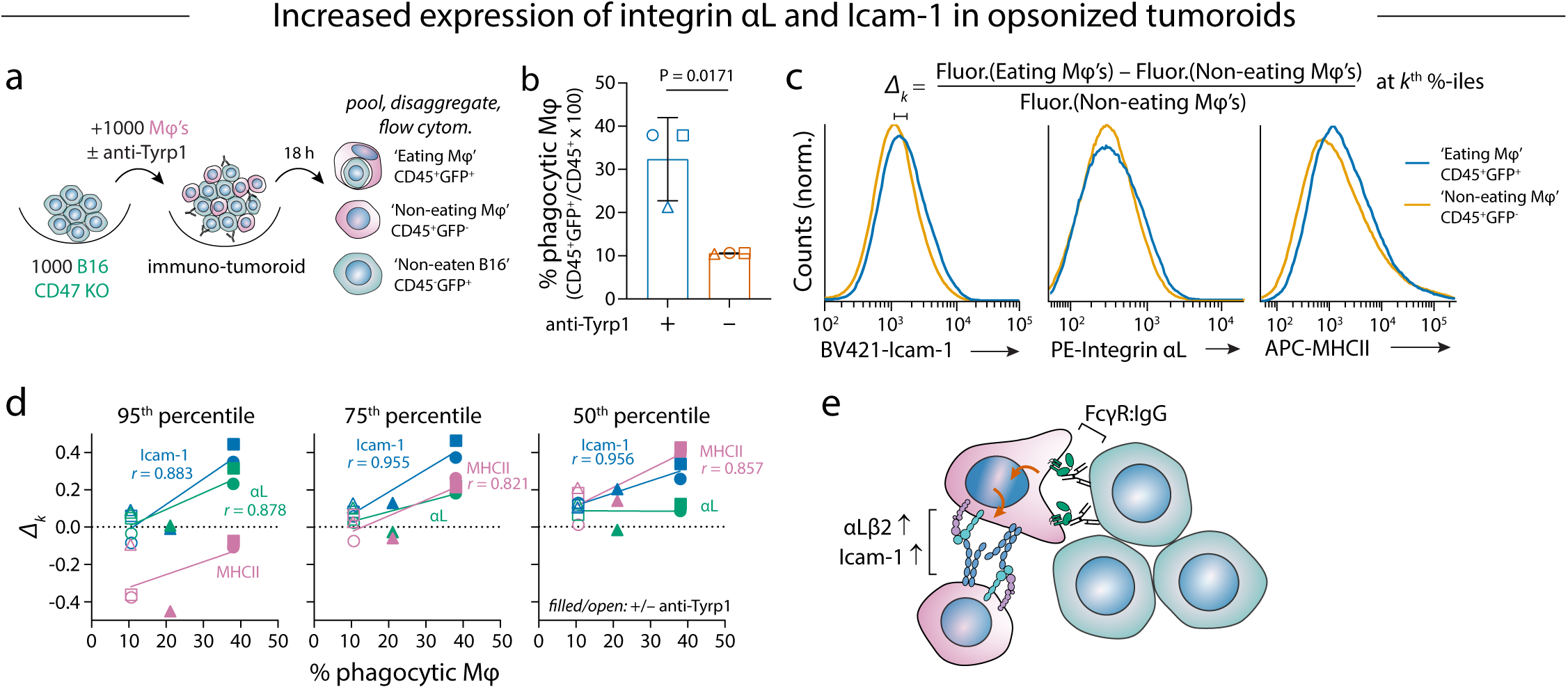
Increased expression of integrin αL and Icam-1 on macrophages from opsonized immuno-tumoroids a. Schematic of tumoroid experiment and disaggregation for flow cytometry staining. b. Flow cytometry analysis quantifying the percentage of phagocytic macrophages (CD45^+^GFP^+^/CD45^+^ x 100%) isolated from immuno-tumoroids opsonized with anti-Tyrp1 or unopsonized. Mean ± s.d., *n* = 3 samples pooled from the same 96-well plate per condition, Student’s *t*-test (unpaired, two-tailed). c. Representative histograms of Icam-1, integrin αL, and MHCII flow cytometry staining of immuno-tumoroid macrophages gated as either ‘eating’ CD45^+^GFP^+^ or ‘non-eating’ CD45^+^GFP^-^. The parameter *Δk* equals the difference between the *k*-th percentile fluorescence intensity value of eating and non-eating macrophages normalized by the value of the non-eating macrophages. d. Correlation between *Δk* at the 95^th^, 75^th^, and 50^th^ percentiles of Icam-1, integrin αL, and MHCII fluorescence (illustrated in panel **c**) and the percent phagocytic macrophages (reported panel **b**). Symbol shapes denote different immuno-tumoroids, and filled and unfilled symbols are opsonized and unopsonized tumoroids, respectively. Pearson coefficients (*r*) are shown for statistically significant correlations (P < 0.05, two-tailed). e. Proposed upregulation of macrophage integrin αLβ2 and Icam-1 downstream of phagocytic signaling networks in immuno-tumoroids based on rightward shift of fluorescence intensity distributions of phagocytic vs. non-phagocytic macrophages.

### Differential Actomyosin Contractility Can Influence Clustering

To determine how macrophage clustering via these receptors might enhance phagocytosis, we next considered their linkage to the actin cytoskeleton via their cytoplasmic domains. Changes in actin organization could conceivably contribute to the differential clustering behavior of macrophage subtypes and would also affect Fcγ receptor-mediated phagocytosis, which is driven by actin polymerization that pushes pseudopod extensions around the target (38). To investigate this possibility, we performed the macrophage clustering assay in the presence of latrunculin A (LatA), an actin polymerization inhibitor, or *para*-amino-blebbistatin (NH2-blebb), an inhibitor of myosin-II-mediated contractility (**Fig.4a,b**). M1 clustering was abrogated in the presence of LatA while the dispersion of M0 macrophages was modestly but significantly decreased (Fig.4a). On the other hand, M0 macrophages were significantly more clustered in the presence of NH2-blebb while clustering of M1 macrophages was unaffected (Fig.4b). Thus, while our results demonstrate a clear requirement for actin filaments in macrophage clustering, they also reveal that clustering is opposed by myosin-II contractility and suggest that myosin activity might be lower in M1 macrophages relative to M0 and M2 macrophages.

To begin to understand which actin structures might be most affected by macrophage polarization, we again analyzed public transcriptomic datasets (Table S1) for genes involved in cytoskeleton organization, adhesion complexes, and mechanosensing, which revealed differential expression of many such genes including *Vcl*, *Actn1*, *Lmna* and others in M1 and M2 BMDMs. In other cell types including mesenchymal stem cells (MSCs), such genes are part of the serum-response factor (SRF) pathway and are downregulated on soft substrates that generally limit cellular contractility (39, 40). Therefore, we specifically examined the expression of actin- and cytoskeleton-associated genes that were previously shown to be differentially expressed in peritoneal macrophages elicited from WT and *Srf* KO mice (41). Many of these SRF-regulated genes are indeed downregulated in M1 or M1’ BMDMs and upregulated in M2 BMDMs after 1 day of polarization, although kinetic profiles suggest more complicated regulation including increased expression within ∼1-2 h in both M1’ and M2 BMDMs (**Fig.4c,** left, **Fig.9a**). We focused on two of the most strongly differentially expressed genes, *Vcl* and *Actn1* that encode the mechanosensitive adhesion protein, vinculin, and the actin crosslinking protein, α-actinin, respectively. In addition to differential expression observed in WT vs. *Srf* KO mouse macrophages (41), both human *VCL* and *ACTN1* have SRF binding peaks in their promoter or intronic regions in the ENCODE ChIP-seq database (**Fig.9b**). Western blotting of lysates from M0, M1, and M2 BMDMs across multiple donor mice revealed differential expression at the protein level, and the correlation between α-actinin and vinculin levels strongly suggests co-regulation (**Fig.4d**, **Fig.10a**). In contrast to M1 BMDMs activated with IFNγ, BMDMs treated with interferon-β from the same microarray dataset do not show downregulation of SRF target genes (**Fig.9c**), which would be consistent with the relative lack of macrophage clustering we observed with IFNα (Fig.2a, Fig.4j,k), another type-I interferon that signals through shared receptors.

SRF regulates its own expression, and *Srf* and its co-factors *Mkl1* and *Mkl2* show slight but generally non-significant downregulation in M1 BMDMs and upregulation in M2 BMDMs after 1 day of polarization (Fig.4c, left) with similar kinetic profiles to *Vcl* and *Actn1* (Fig.9a). In contrast to these genes, genes encoding proteins involved in actin branching including components of the Arp2/3 complex and Was-family members are not significantly affected by macrophage polarization, at least at the 24 h timepoint analyzed here (Fig.4c, right).

The *Lmna* gene encoding the nuclear envelope protein lamin-A and the alternative splicing product lamin-C is among the genes differentially regulated across all the transcriptomic datasets we analyzed, and genes encoding components of the linker of nucleoskeleton and cytoskeleton (LINC) complex (*Sun2*, *Syne2*) are also downregulated in M1’ macrophages (Fig.4c). We previously showed that lamin-A,C knockdown in MSCs feeds back to downregulate SRF target gene expression (39), and *Lmna* was also shown to be downregulated in BMDMs in response to LPS with further regulation of lamin-A,C levels by protein degradation (42). We confirmed differential expression of lamin-A,C among M0, M1, and M2 BMDMs by Western blotting (**Fig.4d**) and proceeded to determine the functional consequences by knocking out lamin-A,C entirely using Cas9^+^ CIM progenitors (**Fig.4e**, **Fig.10b**). Similar to M1 BMDMs, macrophages differentiated from lamin-A,C KO CIMs had decreased levels of vinculin and α-actinin protein and were more clustered on low-adhesion substrates than control macrophages (**Fig.4e,f**, **Fig.10c**). Lamin-A,C KO macrophages also exhibited increased phagocytosis of both IgG-opsonized latex particles and anti-Tyrp1-opsonized B16 cells (**Fig.4g**). These results suggest that lamin-A,C levels regulate SRF target genes in macrophages as observed for MSCs and that this influences macrophage functions including phagocytosis (**Fig.4h**), with similar pathways potentially operating in M1 macrophages.

Based on our transcriptomic analysis and functional studies, we hypothesized that weaker cortical actomyosin in M1 macrophages would pose less of a barrier to the generation of cellular protrusions, including pseudopod protrusions involved in phagocytosis (Fig.4c, center). Addition of blebbistatin to mesenchymal cells often leads to more dendritic-shaped cells (43), consistent with the hypothesis. Likewise, M1 BMDMs appear more dendritic and less spread than M2 BMDMs on rigid glass substrates (Fig.4c). Addition of NH_2_-blebb promotes clustering of M0 BMDMs on low-adhesion surfaces (Fig.4b) but does not impede elimination of the B16 CD47 KO cells from immuno-tumoroids (**Fig.11**). Together, this all suggests that the cytoskeleton organization that favors clustering of macrophages might also be highly conducive to phagocytosis.

### Macrophages extend intrusive pseudopods as ‘intrudopodia’ to disrupt solid tumor adhesions

Adhesions between cancer cells in solid tumors potentially act as physical barriers to the extension of pseudopods from phagocytes, which would thereby limit engulfment, but studies of phagocytosis of adherent or cohesive targets are generally lacking (30). It is conceivable that macrophages first disrupt target cohesion and then begin to engulf detached cells, which we hypothesized might occur through either proteolysis of cell-cell adhesion molecules or myosin-II-dependent pulling forces (**Fig.11a**). However, we observed that immuno-tumoroid clearance proceeds normally in the presence of a protease inhibitor cocktail and in the presence of NH_2_- blebb (**Fig.11b-d**), suggesting that neither mechanism contributes significantly to phagocytic elimination of cohesive solid tumor cells.

To better visualize phagocytosis of cohesive targets, we generated 3D spheroids of B16 CD47 KO cells using growth medium containing methylcellulose and then transferred spheroids onto coverslips for time-lapse imaging over the course of several hours after adding macrophages and anti-Tyrp1 (**Fig.5a**, **Fig.12a-c**). After 1-2 days, spheroids can be eliminated similar to tumoroids (Fig.1b), even though spheroids are more cohesive (and less fractal) than tumoroids. At early time points, macrophages were seen to form small clusters on the spheroid periphery rather than forming large central aggregates per tumoroid clusters (Fig.1c). Macrophage interactions with B16’s included protrusions that wedged between B16’s and also pseudopod-like protrusions that with time disrupted B16 cell-cell adhesions. We term the latter structures intrusive pseudopods or ‘intrudopodia’ and propose that they play key roles in disrupting adhesions among solid tumor cells and thereby facilitate phagocytosis.

Analysis of multiple spheroids reveals that intrudopodia and phagocytosis are relatively common. In some snapshots, two macrophages extend intrudopodia in proximity of one B16 cell that is ultimately dislodged and engulfed by one of the macrophages (Fig.5a, Fig.12c). The overall number of intrudopodia over time indeed exceeded the number of successful phagocytic events by ∼5-fold for each spheroid (Fig.5a, plot). Moreover, given that individual macrophages exhibit no more than 1-2 pseudopodia or intrudopodia at any given time, this overall excess relative to phagocytic events is consistent with multiple intrudopodia cooperating for phagocytosis. Also, some intrudopodia meander into the spheroid but do not lead to productive engulfment (**Fig.12d**).

To determine whether intrudopodia might be relevant to phagocytosis in vivo, we re-examined F4/80-stained sections of B16 CD47 KO lung metastases and focused on regions with aggregated macrophages. We observed macrophages that appeared to form wedges and intrudopodia between B16’s based on their extensions between B16 nuclei (**Fig.5b**). While fixed sections provide only a single time point that does not reveal intrudopod dynamics, this analysis is the first evidence for the extension of cellular protrusions between solid tumor cells that conceivably represent a required step for phagocytosis of such targets. Furthermore, we hypothesize that the ability of macrophage clusters to extend multiple intrudopodia, perhaps in a temporally coordinated and cooperative manner, would lead to more efficient disruption of target adhesions than intrudopodia from an isolated macrophage (**Fig.5c**).

### Anti-tumor macrophages in human tumors express high levels of ITGAL and low levels of actin crosslinking genes

To begin to extend our finding to human cancers, we focused on recently described CS^hi^ anti-tumor macrophages that were defined on the basis of a high ratio of *CXCL9* to *SPP1* gene expression (1). Analysis of other cell-type-gene correlations reported in that study revealed that macrophage expression of *ITGAL* and phagocytosis-related genes (*SLAMF7*, *FCGR3A*) correlate positively with the abundance of CS^hi^ macrophages (**Fig.13a**). On the other hand, macrophage expression of multiple genes encoding actin crosslinking and bundling factors (*FLNB*, *TNS1*, *ACTN4, FSCN1*) were found to be negatively correlated. The positive correlation between *ITGAL* expression and CS^hi^ macrophages was also observed in three other cell types (dendritic cells, B cells, T cells) in addition to macrophages, which is similar to the positive correlation across multiple immune cell types observed for the transcription factor *STAT1* that lies downstream of IFNγ signaling (**Fig.13b**). We therefore analyzed the promoter region of human *ITGAL* by querying ChIP-seq experiments in the ENCODE database. Our analysis revealed the presence of binding peaks for STAT1 as well as the macrophage master regulator transcription factor PU.1 and known *ITGAL* regulators (44) RUNX3 and CEBPβ (**Fig.13c**). Importantly, high expression of *ITGAL* associates with patient survival across many different human cancer types (**Fig.13d**). Although other immune cells including T cells would be expected to contribute to this effect, the trend remains even when the analyses are limited to subsets of patients with high numbers of macrophages or decreased numbers of CD8^+^ T cells. Thus, despite the limitations of the M1/M2 polarization paradigm in capturing the full complexity of macrophage subtypes in tissues, differences in adhesion receptor expression and actin organization that promote clustering of mouse macrophages in our reductionist approaches are also potential features of anti-tumor macrophages in human cancer.

## Discussion

Macrophage clustering is a classic phenomenon and one way in which macrophages might contend with objects that are large and difficult to engulf. Although various groups have made qualitative descriptions of macrophage ‘clusters’, ‘aggregates’, and ‘nests’ in human tumors (1, 17, 19, 20), the mechanisms by which they form and any unique functions that are enabled by this spatial organization are not yet understood. Here, we uncovered an association between macrophage clustering and phagocytosis in tumors and immuno-tumoroids and determined that clustering results from upregulation of specific adhesion receptors on macrophages (Fig.2c-h, Fig.3c-e) and decreased actomyosin contractility (Fig.4b).

Macrophages are notorious for making strong adhesions to cell cultureware, and mouse BMDMs are frequently differentiated on bacteriological Petri dishes rather than tissue culture-treated plastic to facilitate detachment, emphasizing the nontrivial nature of the macrophage clusters reported here that do not adhere to their substrate. Our substrates were made poorly adhesive by multiple methods and contained no specific adhesion ligands beyond what might have physiosorbed from serum in the medium. In some M2 macrophages on low adhesive surfaces, an elongated morphology with lamellipodia-like protrusions and trailing edges suggests they can adhere to the substrate more so than M1 macrophages that preferentially adhere to one another in clusters (Fig.4a). With sufficient time, M2 macrophages could have modified the surfaces and produced their own extracellular matrix such as fibronectin as reported by others (45) to promote adhesion and migration. Across different assay types, multiple timescales of interest emerged ranging from minutes for cluster fluidity in micropipette aspiration (Fig.1h) and intrudopodia formation in spheroids (Fig.5a), to hours for macrophage cluster formation (Fig.2c) and changes in gene expression (Figs.S6,S9), to days for macrophage dispersion following M2 (re)polarization or immuno-tumoroid elimination (Fig.2d, Fig.8c,e). We did observe one example of sexual dimorphism in these assays, namely that BMDMs derived from female donor were slower to disperse in M2 conditions following M1-priming (Fig.2f). It is unclear at present what causes this difference, but the trend is consistent with greater pro-inflammatory cytokine responses in female mice (reviewed in (46)).

Our analysis of differential gene expression among polarized mouse BMDMs and functional studies using gene knockout CIMs implicate integrin αLβ2 binding to Icam-1 in mediating macrophage homotypic cell-cell adhesions in clusters (Fig.2c-h, Fig.3e). Upregulation of integrin αL was previously reported for cultured human monocytes and mouse macrophages treated with either IFNγ or LPS, and a role for this integrin in mediating homotypic cell adhesion of monocytes was proposed based on antibody blockade experiments (47–50). Those monocyte aggregates seem much smaller than the M1 aggregates in our work, but this might reflect differences in substrate adhesivity and geometry (i.e. flat culture dish vs. passivated U-bottom well). Moreover, we did not observe inhibition of M1 clustering with anti-αL IgG or F(ab’)2. Instead, some antibodies appeared to promote aggregation of M0 and M2 macrophages (Fig.7), which we attribute to agglutinating activity that might be harnessed to engineer synthetic multicellular macrophage aggregates in the future. There are likely multiple signaling pathways that can result in upregulation of *ITGAL* in macrophages. In addition to IFNγ signaling and a potential role for STAT1 (Fig.13), STAT5 was shown to mediate increased expression upon LPS stimulation in mouse macrophages (49) while upregulation of integrin αL in T cells following T cell receptor crosslinking was dependent on extracellular-regulated kinase (ERK) signaling (51). The latter is noteworthy because ERK signaling is also implicated in transcriptional changes downstream of Fcγ receptor stimulation on macrophages by either antibody crosslinking (52) or optogenetic methods (27), which might explain our observation of increased integrin αL on phagocytic macrophages isolated from IgG-opsonized immuno-tumoroids (Fig.3c,d). Despite a clear role for integrin αLβ2 and Icam-1 in macrophage clustering, complete knockout of any of the three components of this receptor-ligand pair only decreases clustering by ∼50%, and none of the knockouts tested showed defects in M0 or M2 dispersion on low-adhesion surfaces (Fig.2g,h).

Therefore, roles for other integrins and scavenger receptors (15, 18) in macrophage cell-cell adhesion and cell-substrate adhesion require further investigation.

Beyond adhesion receptors, perturbations to the macrophage cytoskeleton affect M1 clustering and M0 dispersion (Fig.4a,b), but opposing effects of actomyosin contractility on multicellular organization have been reported in different cell types. In mixtures of embryonic germ cells, ectoderm cells with high cortical tension preferentially occupy the interior region of mixed clusters, but this organization is disrupted by myosin-II inhibition with blebbistatin and other perturbations to cellular contractility (23). Breast cancer cell lines in which cell adhesion is mediated by E-cadherin behave similarly, whereas cell lines with integrin-mediated adhesions showed the opposite trend with blebbistatin instead promoting aggregation and sorting to the center of mixtures (24). Macrophages seem to more closely resemble the latter in that myosin-II inhibition of contractility promotes clustering, which may be consistent with integrin-mediated adhesions proposed here. The precise role of myosin-II in this process is unknown, but at least in T cells migrating on surfaces coated with Icam-1, myosin-II contractility opposes adhesion between αLβ2 and Icam-1 to facilitate uropod retraction (53). It is possible that myosin-II activity likewise opposes the αLβ2-Icam-1 interaction between clustering macrophages and that it is also required for migratory processes that lead to dispersion of M0 and M2 macrophages. We infer low actomyosin activity in M1 macrophages from 1) the similar clustering behavior of M1 macrophages and NH_2_- blebb-treated M0 macrophages (Fig.4b) and 2) from gene expression profiles indicating decreased levels of SRF target genes in M1 and M1’ macrophages (Fig.4c). The MRTF family of SRF transcriptional co-activators regulates expression of genes encoding contractile and adhesion complex proteins. Knockdown of the SRF target *LMNA* in human MSCs feeds back on the pathway to downregulate other SRF/MRTF target genes (39, 40), and a similar effect appears to occur in mouse macrophages genetically engineered to knock out lamin-A,C expression. The SRF/MRTF pathway has been implicated previously in regulating some macrophage responses to immune stimuli such as interferons (54), zymosan (55), and LPS (5). Mouse BMDMs activated by LPS and peritoneal macrophages activated by zymosan upregulate SRF target gene expression at early timepoints (∼hours) after stimulation. Our analysis of kinetically resolved macrophage transcriptomes also indicates early upregulation of many SRF targets (*Vcl*, *Actn1*, *Lmna*) upon activation with IFNγ + LPS followed by downregulation by 24 h later (Fig.9a). The early increase in transcription could coincide with a transient (∼2 h), myosin-II-dependent contractile stage recently demonstrated to occur in mouse BMDMs following activation with IFNγ + LPS (56). Nonetheless, longer timescales are of relevance in tumoroids grown for days and especially in tumors.

Various studies have reached different conclusions on whether myosin-II is required for FcγR- mediated phagocytosis, but myosin-II accumulation at the phagocytic synapse and constriction of deformable targets have both been observed and could contribute to engulfment (57, 58). Nonetheless, myosin-II inhibition with either blebbistatin (30) or the more photostable *p*-amino- derivative, NH2-blebb, (Fig.11) did not impede elimination of IgG-opsonized immuno-tumoroids, and we hypothesize here that lower cortical actomyosin contractility in M1 macrophages can facilitate phagocytic pseudopod formation by lowering the barrier for membrane protrusions. Thus, low actomyosin contractility that favors macrophage clustering may promote phagocytosis of cohesive targets in two ways: 1) enhancing pseudopod formation (Fig.4) and 2) increasing the density of macrophages locally to permit coordination between intrusive pseudopods or ‘intrudopodia’ for more effective disruption of target cell adhesions (Fig.5). We consider intrudopodia to be distinct but at least conceptually related to different types of actin-based protrusions described as participating in phagocytosis in *Drosophila* embryos and macrophage phagocytosis of adherent bacteria. Specifically, filopodia-like protrusions and cellular ‘arms’ have been proposed to overcome spatial confinement and enable cooperativity between phagocytic cells, respectively (59, 60). Filopodia and lamellipodia have also been implicated as well in a ‘hook-and-shovel’ mechanism to detach surface-bound *E. coli* for subsequent phagocytosis (61). These structures together with the intrudopodia discovered here reveal the robustness of phagocytosis even in physically challenging environments including solid tumors.

**Figure 4.**
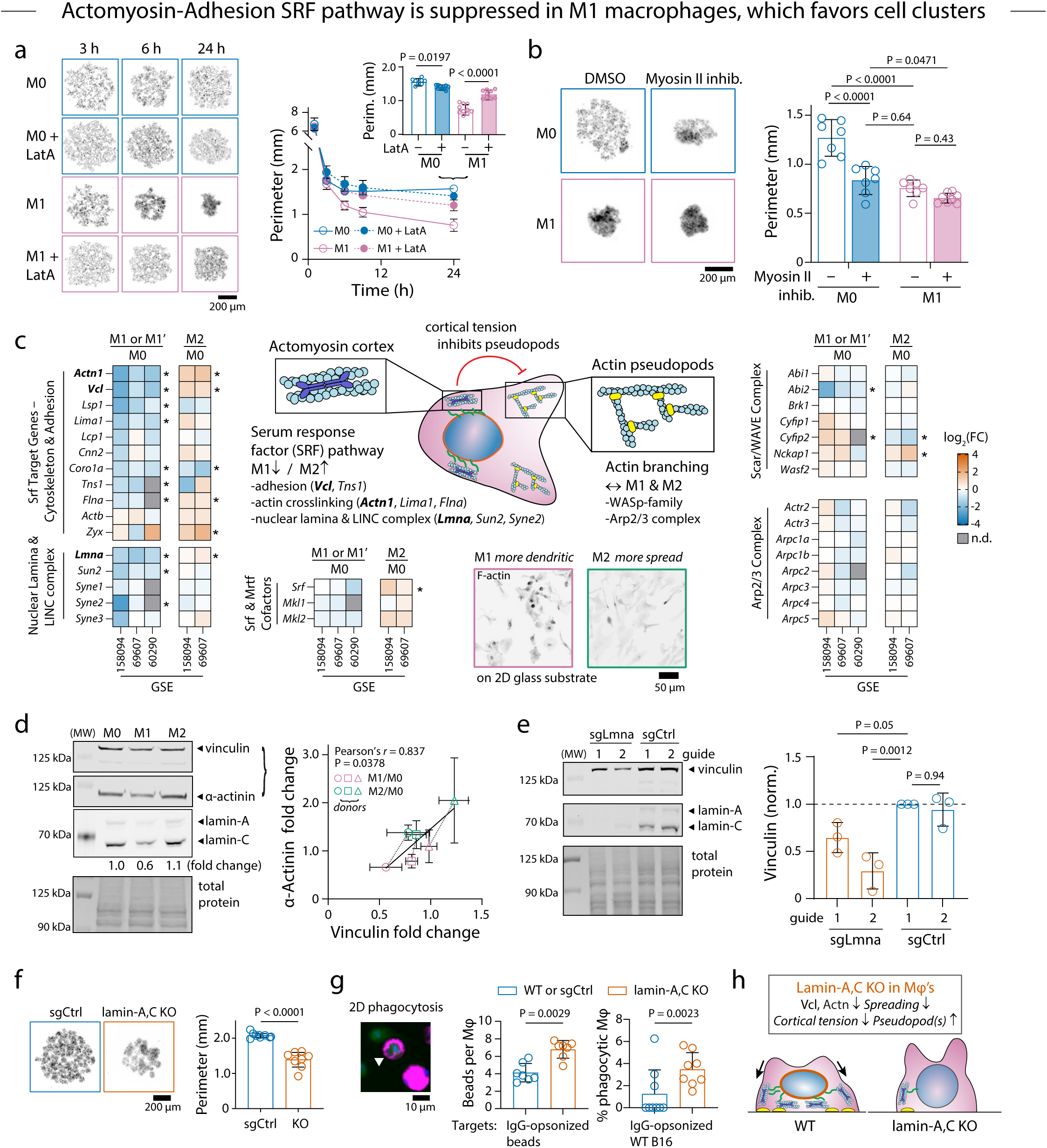
SRF contractile-adhesion pathway is suppressed in M1 macrophages, which favors cell clusters a. Representative fluorescence images and convex hull perimeters of M0 and M1 BMDMs as a function of time on low-adhesion U-bottom well plates with 1 μM latrunculin A (LatA) or DMSO vehicle control. The inset bar graph depicts convex hull perimeters at *t* = 24 h. Mean ± s.d., *n* = 8-10 wells per condition, ordinary two-way ANOVA with Tukey’s multiple comparisons test. Scale bar 200 μm. b. Representative fluorescence images and convex hull perimeters of M0 and M1 BMDMs after 1 day on low-adhesion U-bottom well plates with 50 μM *p*-amino-blebbistatin (myosin-II inhibitor) or DMSO vehicle control. Mean ± s.d., *n* = 7 wells per condition, ordinary two-way ANOVA with Tukey’s multiple comparisons test. Scale bar 200 μm. c. Heat maps of fold changes in expression of (*left*) Srf target genes including actin bundling proteins, adhesion proteins, and nuclear lamina/LINC complex proteins and (*right*) genes encoding actin branching-associated factors between polarized (M1, M1’ or M2) and unpolarized (M0) BMDMs from datasets listed in Table S1. Schematic (*center*) of the hypothesized antagonism between the cortical actomyosin cytoskeleton and pseudopod protrusions driven by branched actin. Fluorescence images M1 and M2 BMDMs stained for F- actin. Scale bar 50 μm. d. Immunoblots of lysates from BMDMs polarized in M0, M1, or M2 medium for 2 days probed with anti-α-actinin, anti-vinculin, or anti-lamin-A,C. The band densities normalized by total protein staining were used to compute M1/M0 and M2/M0 fold changes, which are visualized for α-actinin and vinculin as a scatterplot to emphasize co-regulation as Srf targets. Mean ± s.d., lysates from each donor were loaded in triplicate, Pearson correlation (two-tailed). e. Immunoblots of lysates from macrophages differentiated from lamin-A,C KO and non- targeting control guide CIM progenitors probed with anti-vinculin and anti-lamin-A,C. The densities of vinculin bands were normalized by total protein staining and then by the density of non-targeting control guide 1. Mean ± s.d., *n* = 3 blots, ordinary one-way ANOVA with Dunnett’s multiple comparisons test. f. Representative fluorescence images and convex hull perimeters of macrophages differentiated from lamin-A,C KO and non-targeting control guide CIM progenitors after 1 day on low-adhesion U-bottom well plates. Mean ± s.d., *n* = 8 wells lamin-A,C KO and 9 wells non- targeting control guide, Student’s *t*-test (unpaired, two-tailed). Scale bar 200 μm. g. Quantification of phagocytosis of 7 μm diameter IgG-opsonized polystyrene beads or WT B16 by macrophages differentiated from lamin-A,C KO or WT CIM progenitors. Mean ± s.d., *n* = 7-9 fields of view per condition, Mann-Whitney test, two-tailed. h. Summary of differences between lamin-A,C KO and WT macrophages.

**Figure 5.**
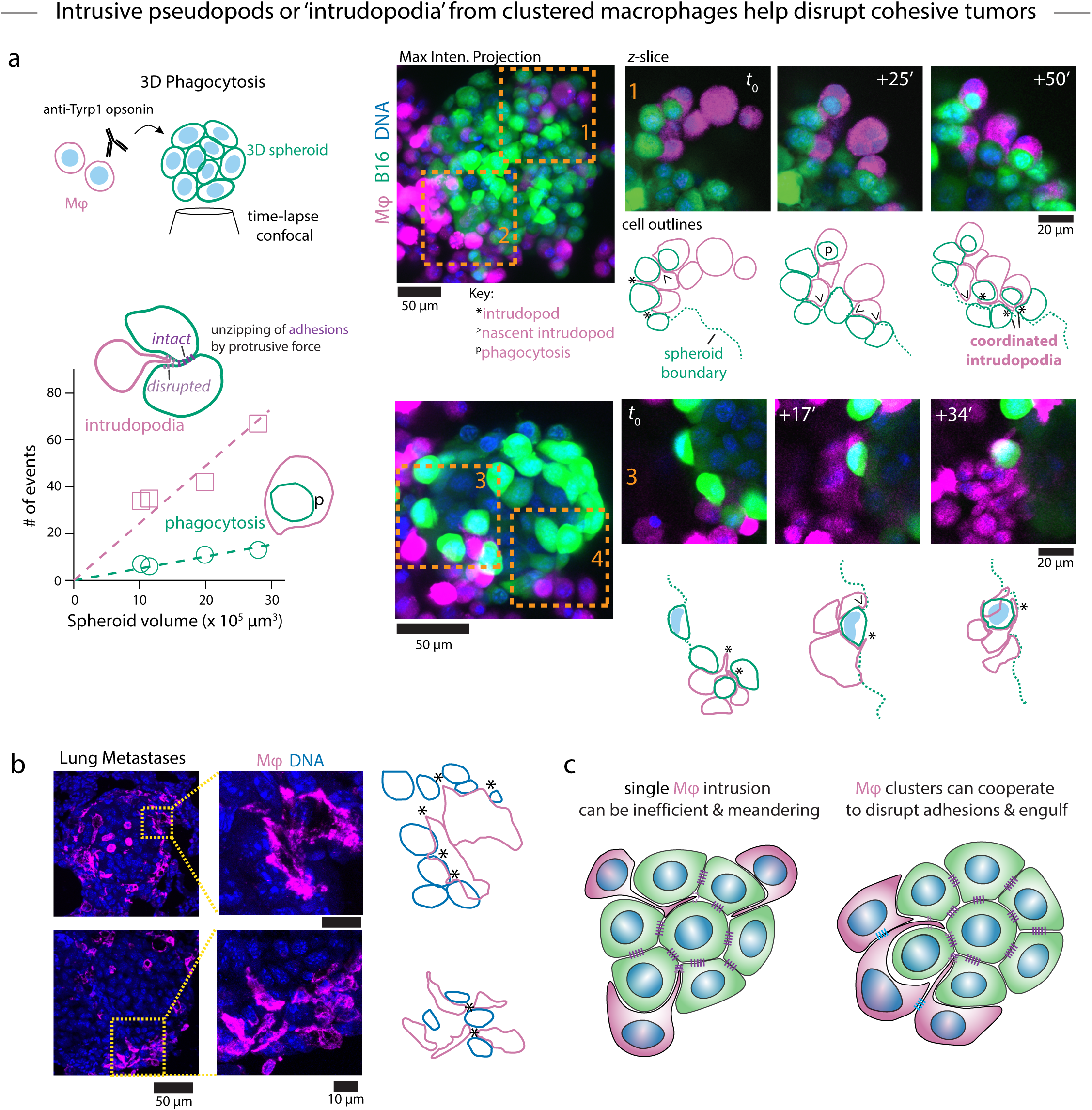
Intrusive pseudopods or ‘intrudopodia’ from clustered macrophages help disrupt cohesive tumors a. Spheroids of B16 CD47 KO cells imaged during phagocytosis. Two projections of confocal stacks of show B16 CD47 KO (GFP^+^, green) to which CellTracker Deep Red-labeled BMDMs and Hoechst 33342 DNA stain were added. Scale bars 50 μm. Zoomed images of a single confocal slice at multiple time points correspond to regions of interest outlined in dotted orange boxes. Scale bars 20 μm. See **Fig.12** for ROI 2 and 4. Cell outlines of B16’s and macrophages in green and magenta, respectively, are shown below the zoomed slices. Events of interest include phagocytosis (p), nascent intrudopodia wedging between B16’s (>), and intrudopodia between B16’s (*). The plot summarizes phagocytic events and intrudopodia observed for *n* = 4 spheroids as a function of the initial spheroid volume, which was calculated from the measured diameter assuming a spherical shape. b. Maximum intensity projection confocal microscopy images of metastatic nodules immunostained for F4/80 (same as Fig.1a). Macrophage cell outlines (magenta) and B16 nuclei outlines (blue) from zoomed regions of interest with putative intrudopodia denoted by *’s. Scale bars 50 μm and 10 μm. c. Hypothesized advantage of clustering macrophages for phagocytosis of cohesive targets.

The disruption of cell adhesions to enable phagocytosis has hitherto unappreciated parallels to the disruption of endothelial junctions during leukocyte diapedesis through blood vessels. At least two routes of diapedesis have been described: the paracellular route through endothelial cell-cell junctions and the transcellular route directly through the cytoplasm of an endothelial cell. Mechanical sensing by leukocytes to identify the route that offers the least mechanical resistance has been proposed based on experiments and theory and referred to as ‘tenertaxis’ (62, 63). Imaging of macrophages invading through the germ band in *Drosophila* embryos further demonstrates how transient disruption of cell adhesions, in this case through cell rounding associated with division, can facilitate macrophage motile processes (64). It is intriguing to speculate that macrophage phagocytosis and trogocytosis (‘cell nibbling’) are analogous to the paracellular and transcellular routes of diapedesis, respectively, and that the predominance of one mechanism over the other depends on sensing physical attributes of the target including size (65), stiffness, and cohesiveness. We do not often observe trogocytic events in immuno-tumoroids (30) or spheroids, but it remains possible that small trogocytosed fragments are not sufficiently resolved in our images. Moreover, trogocytosis might increase in frequency as target cell cohesion increases and phagocytic disruption of adhesions no longer represents the path of least mechanical resistance. The tumoroid and spheroid models should be useful to test this hypothesis in future experiments.

## Methods

### Cell Culture

Knockout (KO) of CD47 in B16-F10 mouse melanoma cells (ATCC CRL-6475) was performed as part of a previous study (66). KO and wild-type (WT) cell lines were cultured in RPMI 1640 medium (Gibco 11835030) supplemented with 10% v/v fetal bovine serum (FBS, Sigma-Aldrich F2442), 1 mM sodium pyruvate (Gibco 11360070) and 1x non-essential amino acid solution (Gibco 11140050). Penicillin and streptomycin (Pen/Strep) were not routinely used when culturing B16 cell lines, although they did not impact results when included.

Bone marrow cells were harvested from femurs and tibias of 8 to 15-week-old male or female C57BL/6J mice (The Jackson Laboratory 000664; RRID:IMSR_JAX:0006664). Red blood cells were depleted with ACK lysis buffer (Gibco A1049201) for 7-8 minutes at room temperature. Cells were cultured in 10 or 15 cm Petri dishes at a density of ∼1 x 10^5^ cm^-2^ for 7 d in differentiation medium comprising Iscove’s modified Dulbecco’s medium (IMDM, Gibco 12440053), 10% v/v FBS, 1% v/v Pen/Strep solution (Gibco 10378016), and 20 ng mL^-1^ macrophage colony-stimulating factor (BioLegend 576404) for 7 d. Additional medium (one-half volume) was added on day 4. Adherent macrophages (referred to as bone marrow-derived macrophages or BMDMs) were used beginning day 7. Typical yields were ∼2.5-3 x 10^7^ BMDMs from ∼5-6 x 10^7^ bone marrow cells per mouse. Conditionally immortalized macrophage (CIM) progenitors originally derived from Rosa26-Cas9 knockin mouse bone marrow cells (33) were maintained as a suspension culture below 5 x 10^5^ mL^-1^ in progenitor medium comprising RPMI 1640 (ATCC Modification, Gibco A1049101), 10% v/v FBS, 1% v/v Pen/Strep, 2 μM β-estradiol (Sigma E2758, dissolved in ethanol at a stock concentration of 10 mM), and 10 ng mL^-1^ granulocyte-macrophage colony-stimulating factor (BioLegend 576304). To differentiate progenitors into adherent macrophages, cells were collected by centrifugation, washed twice with phosphate-buffered saline (PBS, Gibco 14190136) containing 1% v/v FBS, and replated at a density of 3.2 x 10^4^ cm^-2^ in Petri dishes containing differentiation medium. Additional medium (one-half volume) was added on day 4. Differentiated CIMs were used beginning day 7. For polarization studies, adherent BMDMs or differentiated CIMs were detached with 1x TrypLE Express (Gibco 12605010) and re-plated at a density of ∼1 x 10^5^ cm^-2^ in differentiation medium on Petri dishes. After re-adhering overnight, the medium was replaced with fresh differentiation medium containing interferon-γ (IFNγ, BioLegend 575302) or interleukin-4 (IL4, BioLegend 574302) to polarize to M1 and M2 states, respectively. Unless otherwise indicated, IFNγ and IL4 were used at a final concentration of 20 ng mL^-1^ and macrophages were polarized for 48 h. Similar treatments were performed with 20 ng mL^-1^ interferon-α (IFNα, BioLegend 752802) or 100 ng mL^-1^ lipopolysaccharide (LPS, Sigma-Aldrich L4516). All cell cultures were maintained at 37 °C, 5% CO2 in a humidified incubator. B16 CD47 KO and CIM progenitors were negative for mycoplasma.

### CRISPR/Cas9 Gene Knockout in CIM Progenitors

Gene editing in CIM progenitors was performed with minor modifications to a previously published protocol (33) using lentiviral vectors to deliver single guide RNA molecules targeting genes of interest (67). Guide RNA sequences (**Table S2**) were designed with the Broad Institute CRISPick webtool (68), synthesized as pairs of complementary single-stranded oligonucleotides (Integrated DNA Technologies) with overhangs suitable for cloning, phosphorylated at the 5’ ends with T4 polynucleotide kinase (New England Biolabs M0201), annealed to generate a double-stranded insert, and ligated with Quick Ligase (New England Biolabs M2200) into a lentiGuide-puro vector that had been digested with BsmBI (Fermentas FD0454) and dephosphorylated with FastAP (Fermentas EF0654). The lentiGuide-puro plasmid was a gift from Feng Zhang (Addgene plasmid #52963; RRID:Addgene_52963) (67). Lentiviral (LV) particles were prepared by transient transfection of HEK 293T cells with a 4:1:4 molar ratio of lentiGuide-puro plasmids:pVSVG:psPAX2 using TransIt-Lenti Transfection reagent (Mirus Bio 6603). The pVSVG and psPAX2 plasmids were gifts from Bob Weinberg (Addgene plasmid #8454; RRID:Addgene_52963) and Didier Trono (Addgene plasmid #12260; RRID:Addgene_12260), respectively. Approximately 2 mL of supernatant containing LV was collected 48 h post-transfection and filtered through a 0.45 μm polyvinylidene fluoride membrane directly onto 5 x 10^5^ CIM progenitors in 0.5 mL progenitor medium in 6-well, non-TC treated plates. Protamine sulfate (Sigma-Aldrich P4020, prepared as a stock solution at 10 mg mL^-1^ in water and sterile filtered) was added to a final concentration of 10 μg mL^-1^, and the plate was centrifuged at 1000 x *g* for 30 min at 32 °C. Additional progenitor medium was added to a final volume of 5 mL. The transduced CIM progenitors were expanded to 12 mL the following day prior to selection with 12 μg mL^-1^ puromycin (Gibco A1113803) beginning one day later. Lamin-A,C KO CIMs required single-cell cloning by limiting dilution in 96- well plates. All other lines were maintained as polyclonal populations.

### B16 CD47 KO Lung Metastasis Model

Lung metastases were established and treated with intravenous anti-Tyrp1 as described (29). Briefly, 2 x 10^5^ B16 CD47 KO cells were injected in a volume of 0.1 mL sterile PBS intravenously via the lateral tail vein of 6 to 12-week-old male C57BL/6J mice. Mice were randomized into treatment and control groups, and mice in the treated group received 250 μg anti-Tyrp1 (BioXCell BE0151) diluted to a final volume of 0.1 mL in PBS on days 4, 5, 7, 9, 13, and 15. The mice were euthanized on day 16, and the lungs were harvested, fixed in 4% paraformaldehyde (Thermo Fisher Chemicals J19943.K2) for 18 h, and transferred to 70% ethanol. Paraffin embedding, sectioning, and immunofluorescence staining were performed by the Molecular Pathology and Imaging Core (RRID: SCR_022420) of the Perelman School of Medicine of the University of Pennsylvania. Immunofluorescence images of F4/80-stained lung sections were captured on a Leica SP8 laser scanning confocal microscope equipped with a 63x/1.4 NA HC PL APO CS2 oil immersion objective or on an Olympus IX-71 epifluorescence microscope equipped with a 40x/0.6 NA LUCPlanFLN objective, pE300-Ultra illumination (CoolLED), and a Prime sCMOS camera (Teledyne Photometrics) controlled by μManager software (69). Macrophages were counted by a blinded observer and assigned to clusters containing *N* = 1, 2, 3… macrophages to obtain a probability distribution *PN* (= *nN*/*n*total) where *nN* is the number of clusters of size *N* and *n*total is the total number of clusters. The number average cluster size, *N*avg, was computed as

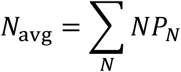

and the size (*N*)-weighted average cluster size, *W*avg, was computed as the ratio of the first and second moments of the cluster size distribution:

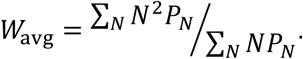

### Tumoroid Assays and Macrophage Aggregation Assays

Low-adhesion surfaces were prepared by two methods. First, as previously described (30), non-TC treated 96-well U-bottom plates were passivated with 100 μL anti-adherence rinsing solution (AAR, StemCell Technologies 07010) for 15 min at room temperature. Alternatively, 50 μL of 30 mg mL^-1^ poly(2- hydroxyethylmethacrylate) (pHEMA, Santa Cruz Biotechnology sc-253284) dissolved in 95% ethanol/5% water was added per well and the solvent was allowed to slowly evaporate with the lid on for 2-3 d. Following either treatment, the wells were washed with PBS and blocked with 1% w/v bovine serum albumin (BSA) in PBS at 37 °C for 1 h. Both treatments generally gave similar results, but we found the polyHEMA treatment to be less variable between wells and between different plate manufacturers or different lots of plates from the same manufacturer.

For immuno-tumoroid assays, B16 cells were prepared as single-cell suspensions in RPMI growth medium at a density of 1 x 10^3^ or 1 x 10^4^ mL^-1^ depending on the experiment, and 100 μL was added per well to previously prepared low-adhesion 96-well plates. The next day, BMDMs were detached from Petri dishes with 1x TrypLE Express and labeled with amine-reactive CellTracker Deep Red dye (Invitrogen C34565). Briefly, macrophages suspended in PBS were labeled with 1 μM of dye from a stock solution in DMSO. Suspensions were incubated at 37 °C for 20 minutes, centrifuged at 300 x *g* for 5 min, and washed with growth medium. In some experiments, BMDMs were primed for 48 h in differentiation medium containing IFNγ or IL-4 prior to use in immuno-tumoroid assays. Macrophage suspensions were adjusted to densities between 1.5 x 10^3^ and 5 x 10^4^ mL^-1^ in RPMI growth medium containing 120 ng mL^-1^ M-CSF. For opsonized tumoroids, 120 μg mL^-1^ anti-Tyrp1 was added to the macrophage suspension. In some experiments, the myosin-II inhibitor *para*-amino-blebbistatin (NH2-blebb, Cayman Chemical 22699) or a protease inhibitor cocktail (Sigma P1860) were added together with the macrophages and anti-Tyrp1 for a final concentration of 20 μM and 0.5% v/v, respectively. After acquiring images of B16 tumoroids corresponding to *t* = 0, 20 μL of the macrophage suspension was added to each well and the plates were incubated at 37 °C for up to four days.

For macrophage clustering assays, BMDMs and differentiated CIMs were detached from Petri dishes with TrypLE Express and labeled with CellTracker Deep Red or Vybrant carboxy- fluorescein diacetate succinimidyl ester Cell Tracer (Invitrogen V12883) as described above. The cell density was adjusted to 1 x 10^4^ mL^-1^ in RPMI growth medium containing 20 ng mL^-1^ M-CSF, and 100 μL was added to each well of previously prepared low-adhesion 96-well plates. Cell suspensions and other solutions were passed through 40 μm nylon cell strainers (Falcon 352340) to remove any particulate debris.

Fluorescence microscopy was performed on the Olympus IX-71 microscope using 4x/0.13 NA UPlanFL, 10x/0.3 NA UPlanFL, and 20x/0.4 NA LC Ach objectives. Images were analyzed in Fiji/ImageJ (70) by applying thresholding (MaxEntropy, Li, or IJ_IsoData algorithms depending on the fluorescence signal) to segment fluorescent cells from background followed by the built-in Convex Hull function on the selected segmented objects. In some cases, manual adjustments to the segmented objects were required. To determine the radial fluorescence profiles, a circle was drawn around each tumoroid as the input for the Radial Profile Angle plugin. The output is the average fluorescence intensity at a radius of *r* pixels, which was then normalized by the total average fluorescence intensity from *r* = 0 to *r*max to approximate the probability distributions *p*(*r*) for M1 and M2 macrophages in co-culture mixtures.

### Drug, Conditioned Medium, and Antibody Treatments

The actin polymerization inhibitor Latrunculin A (Lat A, Cayman Chemical 10010630) was prepared as a stock solution at 1 mM in DMSO and used at a final concentration of 1 μM. NH2-blebb was prepared as a stock solution at 20 mM in DMSO and used at a final concentration of 20 or 50 μM. Cycloheximide (CHX, MP Biomedicals 194527) was prepared as a stock solution of 35 mM in DMSO and used at a final concentration of 35 μM. Pertussis toxin from *Bordetella pertussis* (Calbiochem 516560) was reconstituted at a concentration of 100 μg mL^-1^ in sterile water and used at a final concentration of 10 ng mL^-1^. All drugs were added to macrophage suspensions immediately prior to seeding in low-adhesion well plates.

For conditioned medium experiments, primary mouse T cells were isolated from the spleens of C57BL/6J mice. Splenocytes were collected by mechanically disrupting spleens and filtering the resulting suspension through a 70 μm nylon cell strainer with RMPI 1640 growth medium. CD4^+^ and CD8^+^ T cells were enriched by negative depletion using MojoSort magnetic isolation beads following the manufacturer’s protocol (BioLegend 480005 and 480007). The enriched cell populations were cultured in a 48-well tissue culture plate at a density of 1 x 10^6^ per 350 μL RPMI 1640 growth medium. To activate T cells, concanavalin A (eBioscience 00-4978-93) was added to a final concentration of 2.5 μg mL^-1^ for 48 h. The conditioned media from T cell cultures were collected, clarified by centrifugation, and mixed with an equal volume of fresh growth medium containing 40 ng mL^-1^ M-CSF for macrophage assays.

For antibody blockade studies, functional grade antibodies (Table S3) were added to macrophage suspensions just prior to addition to low-adhesion surfaces. F(ab’)2 fragments were prepared from anti-CD11b clone M1/70, anti-CD18 clone M18/2, and rat IgG2a isotype control clone RTK2758 antibodies by digestion on immobilized pepsin (Pierce 20343) for 4-6 h at 37 °C in 0.1 M sodium acetate buffer, pH ∼3.8. The digest reactions were collected, and the Fc fragments were removed using a 50 kDa MWCO column (Amicon UFC5050). The final products were buffer exchanged to PBS, analyzed by SDS-PAGE and UV/Vis spectroscopy to determine the purity and concentration, and sterilized with a 0.22 μm Spin-X centrifuge filter tube (Corning 8160).

### Micropipette Aspiration

Micropipette aspiration experiments were performed as previously described (30). Glass capillaries (1 mm outer diameter/0.75 mm inner diameter, World Precision Instruments TW100-3) were pulled on a Flaming/Brown type pipette puller (Sutter Instruments P- 97), scored with a ceramic tile, broken to a diameter of approximately 50 µm, and bent by heating the tapered region on a de Fonbrune microforge. The micropipette was backfilled with 3% w/v BSA in PBS and connected to a dual-stage water manometer. Macrophage or B16 cell clusters were transferred using a wide-bore pipette tip from 96-well plates to a glass-bottom dish containing growth medium. Aspiration was applied manually with a 0.5 mL syringe as the aspiration pressure difference was measured with a calibrated DP-15 pressure transducer (Validyne). Time-lapse brightfield microscopy of aspiration was performed on a Nikon TE300 microscope with a 20x/0.5 NA Plan Fluor objective and images were acquired on an Evolve Delta EMCCD camera (Teledyne Photometrics) using μManager software.

### Flow Cytometry

BMDMs and differentiated CIMs were detached from Petri dishes with 1x TrypLE Express for 15-20 min at 37 °C. Cell suspensions were stained with 1:500 Zombie yellow viability dye (BioLegend 423103) in PBS for 15 min at room temperature in the dark and washed twice with FACS buffer (1% w/v BSA and 0.1% w/v sodium azide in PBS). To block Fc receptors, 1 μg mL^-1^ anti-CD16/CD32 (clone 2.4G2, BioXCell BE0307) and 0.25 μg mL^-1^ anti-CD16.2 (clone 9E9, BioLegend 149502) were added to cell suspensions containing ∼1 x 10^7^ mL^-1^ in FACS buffer for 10 min on ice. Primary antibodies listed in Table S3 were added and suspensions were incubated for 30 min on ice. After washing three times with FACS buffer, cells were fixed in FluoroFix buffer (BioLegend 422101) for 1 h at room temperature, washed with FACS buffer, and stored at 4 °C until analysis on a BD LSR II flow cytometer. For flow cytometry analysis of immuno-tumoroids, cells were collected 18 h after addition of BMDMs, and the contents of all wells from a single 96-well plate were pooled as one replicate. The cells were dispersed to single-cell suspensions, washed with FACS buffer, incubated in FACS buffer containing Fc blocking antibodies, and stained with fluorophore-conjugated antibodies against CD45, MHCII, integrin αL, and Icam-1 (Table S3). Samples were analyzed on a BD Symphony A3 lite cytometer. Data were analyzed with FCS Express 7 (De Novo). Flow cytometry data appearing in the same plot were acquired at the same time, or the cytometer photomultiplier gains were adjusted using UltraRainbow calibration beads (Spherotech URCP-38-2K).

### Immunofluorescence and F-actin Staining

Differentiated BMDMs were detached from Petri dishes and re-plated on glass coverslips at a density of ∼5.5 × 10^3^ cm^-2^ in differentiation medium. IFNγ or IL4 were added after cells attached to the glass. Two days later, the BMDMs were washed with PBS, fixed with 4% w/v formaldehyde (diluted in PBS from 16% w/v stock, Pierce 28908), permeabilized with 0.2% v/v Triton X-100 in PBS, and stained with 1:40 (∼165 nM) rhodamine phalloidin in 5% v/v normal donkey serum (Sigma-Aldrich D9663) in PBS. WT or lamin-A,C KO CIM progenitors were differentiated directly on glass coverslips for 7 d and then fixed and permeabilized with ice-cold methanol for 15 min at 4 °C and blocked in PBS containing 5% v/v normal donkey serum, 0.3% v/v Triton X-100, 10 μg mL^-1^ anti-mouse CD16/CD32, and 2.5 μg mL^-1^ anti-mouse CD16.2 for 1 h at room temperature. Coverslips were incubated with anti-lamin-A,C in blocking buffer for 18 h at 4 °C, washed, and incubated with 10 μg mL^-1^ Alexa Fluor Plus 647- conjugated donkey anti-mouse IgG [H+L] and 0.2 μg mL^-1^ DAPI (Sigma-Aldrich D9542) for 1 h at room temperature, washed, and mounted on glass slides with ProLong Gold Antifade (Invitrogen P36930). Imaging was performed on the Olympus IX-71 with a 60x/1.25 NA UPlanFL objective. To quantify lamin-A,C nuclear staining in macrophages, DAPI-stained nuclei were segmented using the Otsu thresholding and Watershed algorithms in FIJI to generate masks, and the lamin- A,C immunofluorescence was integrated over each nucleus.

### Western Blotting

Cell lysates were prepared in radioimmunoprecipitation assay (RIPA) buffer (Cell Signaling Technologies 9806) supplemented immediately before use with a protease inhibitor cocktail (Sigma P8340) and a phosphatase inhibitor cocktail (Sigma P0044). Lysis was completed by multiple freeze-thaw cycles and the solution was sheared by passing through a 27g needle several times before clarification by centrifugation at 10,000 *g* for 30 min at 4 °C. The protein concentration was determined by BCA assay (Pierce 23235) and adjusted to load 10 μg of protein per lane in sample buffer (Invitrogen NP0007 or LC2570) containing 50 mM dithiothreitol. Proteins were denatured and reduced by heating at 75 °C for 10 min prior to separation by gel electrophoresis in 15-well, 1 mm thick Bolt Bis-Tris Plus 4-12% gradient polyacrylamide gels (Invitrogen NW04125) in MOPS buffer at 200 V for approximately 30 min. Transfer to a nitrocellulose membrane (Invitrogen IB301002) was performed with the iBlot dry transfer system. Membranes were air dried for 1 h, then rehydrated and stained with Revert700 Total Protein Stain (LiCor 926-11011) according to the manufacturer’s instructions. Stained membranes were imaged on the Odyssey M imager (LiCor) to normalize for protein loading and then destained for 3 min with 0.1 M sodium hydroxide, 30% methanol solution. Destained membranes were blocked with 5% w/v BSA in PBS for 1 h at room temperature with agitation. Primary and secondary antibodies were diluted in 5% w/v BSA in PBS with 0.2% v/v Tween (PBST). Membranes were incubated with primary antibody (Table S3) solution overnight at 4 °C with agitation, washed three times with PBST for 10 min, incubated with secondary antibody solution for 1 h at room temperature, and washed three times with PBST for 10 min. Fluorophore-conjugated secondary antibodies were detected on the Odyssey M imager. All quantitation of Western blots was performed with Empiria Studio Software (LiCor).

### 2D Phagocytosis

Differentiated lamin-A,C knockout and guide control or WT CIMs were seeded in a 24-well tissue culture plate at a density of 5.3 x 10^4^ cm^-2^ in 1 mL differentiation media and allowed to attach overnight. CIMs were labeled with CellTracker Deep Red. For bead phagocytosis assays, streptavidin-coated polystyrene particles (∼7 μm diameter, Spherotech SVP-60-5) were fluorescently labeled with 1 μM Atto550-biotin (Sigma-Aldrich 28923) in PBS for 20 min at room temperature, washed, and opsonized with rabbit anti-streptavidin antiserum diluted 1:400 in serum-free medium (SFM) comprising IMDM supplemented with 0.5% w/v BSA.

Labeled and opsonized beads were added to macrophages at a 5:1 ratio in 1 mL SFM and the plate was incubated at 37 °C for 90 min. The wells were then washed with PBS, fixed in 4% paraformaldehyde for 30 min, blocked in 5% v/v normal donkey serum in PBS for 1 h, and stained with 2 μg mL^-1^ Alexa Fluor Plus 647-conjugated donkey anti-rabbit IgG [H+L] and 2 μg mL^-1^ Hoechst 33342 dye (Invitrogen 62249) for 1 h to label non-internalized beads and DNA, respectively. For B16 phagocytosis assays, WT B16 were opsonized with 20 μg mL^-1^ anti-Tyrp1 in SFM for 30 min on ice, washed with SFM, and added to macrophages a 2:1 ratio in 1 mL SFM. After 90 min at 37 °C, the wells were also washed with PBS, fixed, and counterstained with Hoechst 33342. Images were acquired on the Olympus IX-71 microscope.

### 3D Spheroid Phagocytosis

To generate spheroids, B16 CD47 KO cells were cultured in RPMI-based growth medium and seeded into non-adherent 96-well U-bottom plates pre-treated with 1% F-127 pluronic (Sigma-Aldrich P2443) solution at a density of 50 cells per well. After 24 h, the culture medium was replaced with RPMI growth medium containing 1% w/v methylcellulose (Thermo Scientific Chemicals 428430500) to promote spheroid formation. Spheroid formation was monitored, and 24 hours after the addition of methylcellulose, spheroids were washed with PBS to remove methylcellulose and moved to a 384-well plate amenable to time-lapse microscopy. The spheroids were then opsonized with 20 µg mL^-1^ anti-Tyrp1, and BMDMs were added to each well at a 5:1 BMDM:B16 ratio.

Time-lapse imaging of spheroids and BMDMs was performed using a Zeiss LSM 880 laser scanning confocal microscope equipped with a LD C-Apochromat 40x/1.1 NA water immersion objective and an environmental chamber maintaining the atmosphere at 37 °C and 5% CO2. Imaging was initiated immediately following the addition of BMDMs, with time-lapse images captured every ∼20 minutes for up to 8 hours. The imaging parameters, including laser settings and acquisition conditions, were optimized to minimize photobleaching and maintain cellular viability throughout the experiment.

### Data Analysis, Statistics, and Curve Fitting

Gene expression in polarized BMDM subtypes was analyzed from the public RNAseq and microarray datasets listed in Table S1. Microarray datasets were analyzed with the GEO2R webtool (https://www.ncbi.nlm.nih.gov/geo/geo2r/) using default options including adjustment of P-values with the false discovery rate method of Benjamani and Hochberg. RNAseq expression data was analyzed by downloading data directly from ref (37) and also by downloading pseudo-aligned transcript counts (mouse_gene_v2.4.h5) from the ARCHS^4^ database (71) followed by processing using Bioconductor packages *edgeR* and *limma* in RStudio 2023.06.0+421 to assess differential gene expression (72, 73). Transcription factor binding to the *ITGAL* promoter region (hg38 chr16:30,470,992-30,472,742) and *SRF*, *VCL*, *ACTN1*, and *LMNA* genes was assessed using the UCSC Genome Browser (http://genome.ucsc.edu) TF ChiP track with data sourced from ENCODE 3 (74). The reported ChIP-seq score was computed by the UCSC Genome Browser project and has a maximum value of 1000. Cell-type-gene correlation coefficients and hazard ratios for select cancer types were reported in ref (1). Survival analysis of patients based on the ratio of *ITGAL* (or *CXCL9*) to *SPP1* was performed using the Kaplan-Meier plotter pan-cancer RNA-seq webtool (kmplot.com) (75). All other statistical analyses and curve fitting were performed in GraphPad Prism 10. Details for each analysis are provided in the figure legends.

## Acknowledgements

We thank members of the Discher Lab for discussion and critical feedback, especially Irena Ivanovska and Karanvir Saini. This work was supported by funding from the National Institutes of Health (NIH) National Cancer Institute (NCI) through grants U01 CA254886 and P01 CA265794 and by the Center for Engineering MechanoBiology (CEMB), an NSF Science and Technology Center, under grant agreement CMMI 15-48571. We acknowledge the following University of Pennsylvania Perelman School of Medicine core facilities and their funding sources. Tissue processing, sectioning, and immunostaining was performed by the Molecular Pathology & Imaging Core (RRID:SCR_022420), which is supported by the Center for Molecular Studies in Digestive and Liver Diseases NIH/NIDDK Grant P30 DK050306. Confocal microscopy was performed in the Cell & Developmental Biology Microscopy Core Facility (RRID:SCR_022373). Flow cytometry data was generated on instruments maintained by the Penn Cytomics and Cell Sorting Shared Research Laboratory (RRid:SCR_022376), which is supported by the Abramson Cancer Center NIH/NCI Grant P30 CA016520.

## Supplementary Figures 1-13 Legends

**Figure S1.**
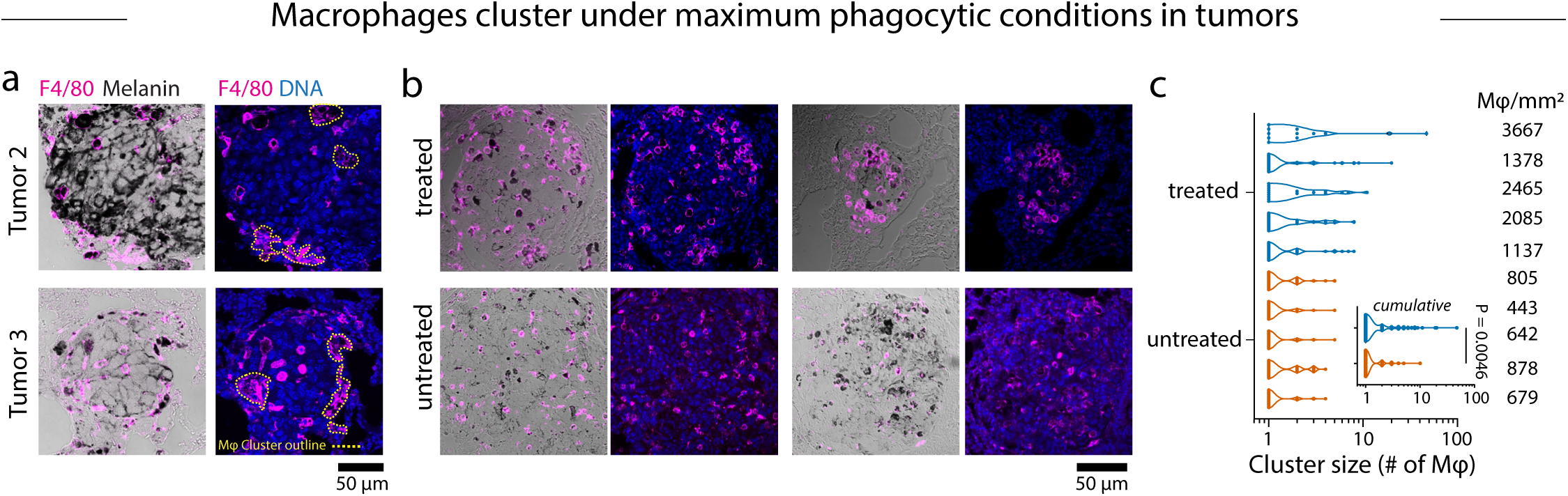
Macrophage clustering in tumors a. Bright-field and confocal fluorescence microscopy maximum intensity projections of two additional metastatic nodules immunostained for F4/80 from mice treated with anti-Tyrp1. The yellow dotted outlines denote macrophage clusters. Scale bar 50 μm. b,c. Representative bright-field and fluorescence images of B16 nodules immunostained for F4/80 (**b**) and cluster size distributions and macrophage density for five nodules each from treated and untreated mice (**c**). Inset: Cumulative cluster size distributions combining clusters from treated and untreated nodules. Kolmogorov-Smirnov test. Scale bar 50 μm.

**Figure S2.**
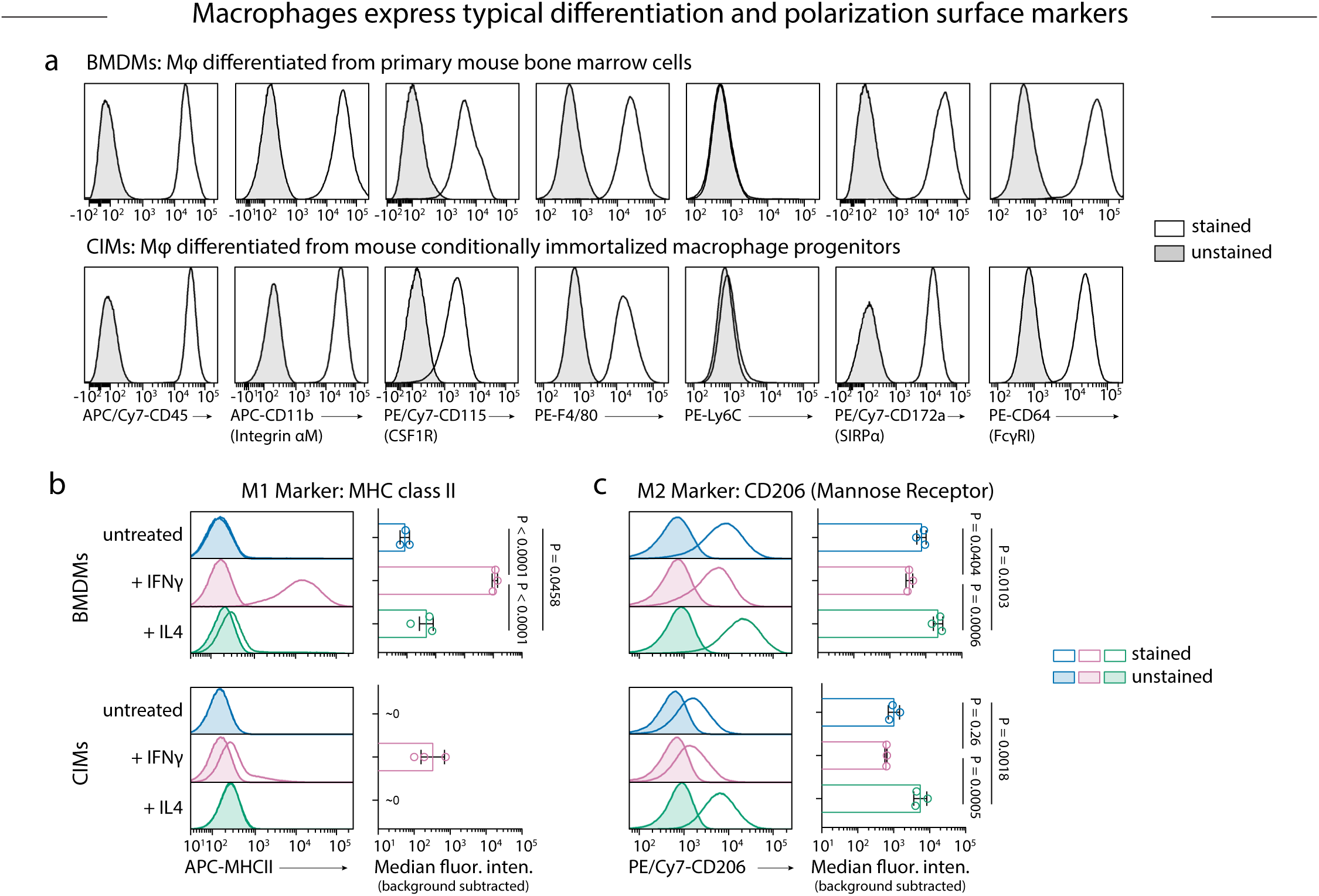
Differentiation and polarization marker expression on BMDMs and CIMs a. Flow cytometry histograms for macrophages differentiated from mouse bone marrow cells (BMDMs, *top*) and conditionally immortalized macrophage progenitors (CIMs, *bottom*) stained with myeloid and macrophage lineage markers: CD45, CD11b, CD115, F4/80, Ly6C, SIRPα, and FcγRI. Filled and unfilled histograms are unstained and antibody-stained samples, respectively. b,c. Flow cytometry staining of untreated, IFNγ-treated, and IL4-treated BMDMs and CIMS for MHCII (**b**) and CD206 (**c**). Histograms show staining of a representative BMDM or differentiated CIM progenitors culture and bar graphs report the median fluorescence intensity (MFI) of stained cells corrected by subtracting the MFI of the unstained culture. Mean ± s.d., *n* = 3 mice for BMDMs or 3 unique cultures of differentiated CIMs, ordinary one-way ANOVA and Tukey’s multiple comparisons test on log-transformed MFI values.

**Figure S3.**
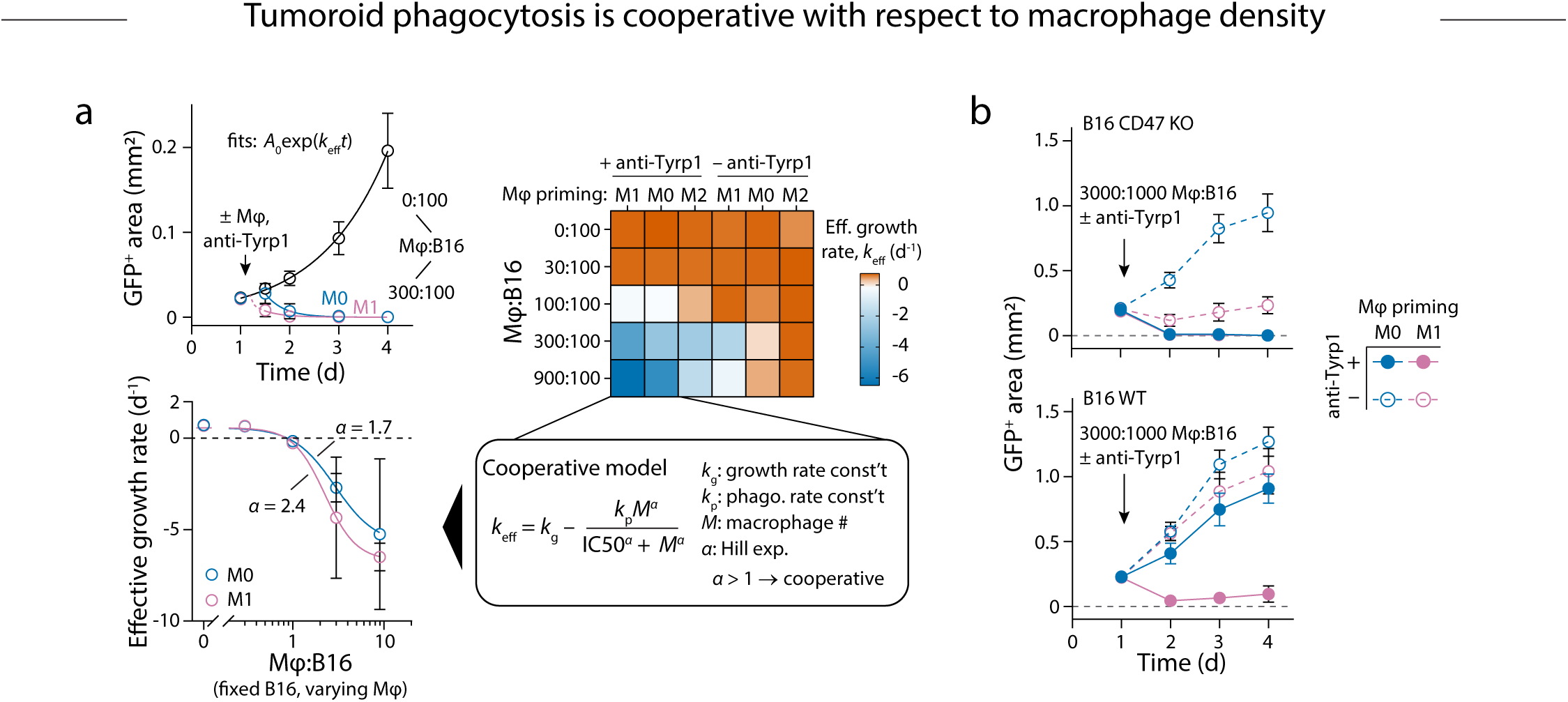
Modeling growth and cooperative phagocytosis in immuno-tumoroids treated with cytokine-primed BMDMs a. Representative growth curves (GFP^+^ projected area vs. time) for tumoroids made from 100 B16 CD47 KO cells and treated on day 1 with 300 M0- or M1-primed BMDMs and 20 μg mL^-1^ anti-Tyrp1 or with anti-Tyrp1 only. The solid lines are fits of the exponential growth equation starting at *t* = 12 h, the first measurement after adding BMDMs, and the dashed line is the extrapolation of the fitted M1 exponential curve to intersect the growth curve of the tumoroid without BMDMs, indicative of a delay time. The heat map depicts the effective growth rate, *k*eff, across different BMDM:B16 ratios and different macrophage priming conditions with or without anti-Tyrp1. The effective growth rates of opsonized tumoroids with M0- or M1-primed BMDMs were fit to the indicated model of exponential B16 growth opposed by Hill-like cooperative phagocytosis with the fitted Hill exponents *α* > 1. Mean ± s.d. for GFP^+^ area or fitted value ± standard error (s.e.) for *k*eff, *n* = 6-8 wells per condition. g. Growth curves for tumoroids made from 1000 CD47 KO (*left*) or WT (*right*) B16 and treated on day 1 with 3000 M0 or M1 BMDMs with or without anti-Tyrp1. Mean ± s.d., *n* = 7-8 wells per condition.

**Figure S4.**
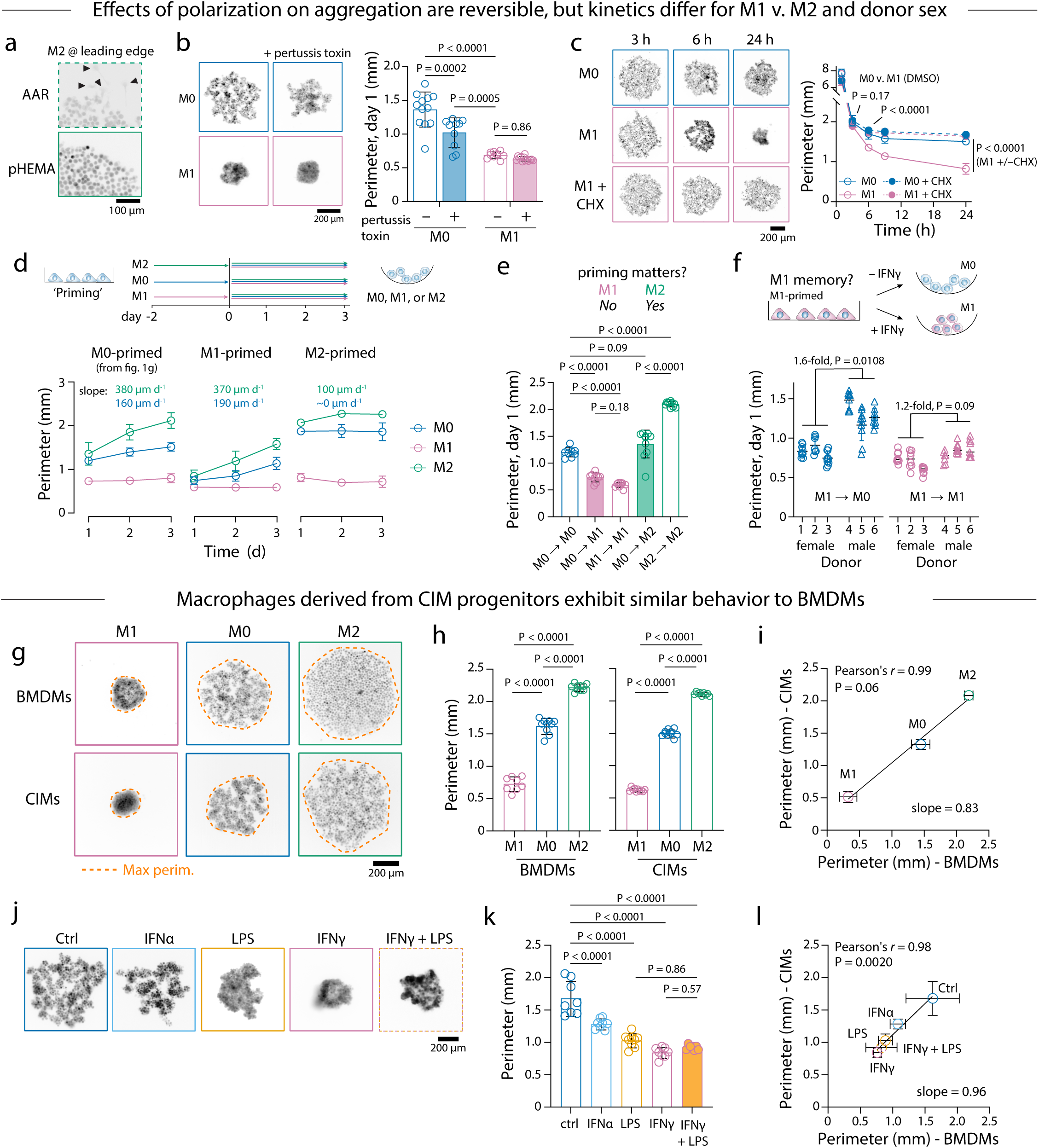
Effects of polarization are reversible, but kinetics differ for M1 vs. M2 and donor sex; macrophages from conditionally immortalized myeloid progenitors (CIMs) show similar behavior to BMDMs a. M2 macrophages in U-bottom well that were treated with anti-adherence rinsing (AAR) solution or coated with poly(2-hydroxyethylmethacrylate) (pHEMA). Arrowheads denote prominent lamellipodia and front-rear polarization in macrophages at the leading edge on the AAR-treated surface. Scale bar 100 μm. b. Representative fluorescence images and convex hull perimeters of M0 and M1 BMDMs after 1 day in low-adhesion U-bottom well plates with or without 10 ng mL^-1^ pertussis toxin. Mean ± s.d., *n* = 10-13 wells per condition, ordinary two-way ANOVA with Tukey’s multiple comparisons test. Scale bar 200 μm. c. Representative fluorescence images and convex hull perimeters of M0 and M1 BMDMs as a function of time on low-adhesion U-bottom well plates with 35 μg mL^-1^ cycloheximide (CHX) or DMSO vehicle control. Mean ± s.d., *n* = 8-10 wells per condition, ordinary two-way ANOVA with Tukey’s multiple comparisons test at the times indicated. Scale bar 200 μm. d. Convex hull perimeters as a function of time after repolarization of BMDMs primed for 2 days in standard 2D culture prior to transfer to low-adhesion U-bottom well plates with M0, M1, or M2 medium. Mean ± s.d., *n* = 8-10 wells per condition. e. Convex hull perimeters of repolarized BMDMs on day 1 for selected transition conditions from d. Mean ± s.d., *n* = 8-10 wells per condition, ordinary one-way ANOVA and Tukey’s multiple comparisons test. f. Convex hull perimeters of BMDMs on day 1 following M1 priming and transfer to low-adhesion U-bottom well plates with M0 or M1 medium. Mean ± s.d., *n* = 8-10 wells for 3 female (circles) and 3 male donors (triangles), nested t-test (two-tailed). f,h. Representative fluorescence images (**g**) and convex hull perimeters (**h**) of BMDMs and differentiated CIMs primed under M0, M1, or M2 conditions for 2 days in 2D culture and transferred to low-adhesion U-bottom well plates for 1 day in M0, M1, or M2 medium, respectively. Mean ± s.d., *n* = 8-10 wells per condition, ordinary one-way ANOVA and Tukey’s multiple comparisons test. Scale bar 200 μm. i. Pearson correlation between the convex hull perimeters of CIMs and BMDMs from **h**. j,k. Representative fluorescence images (**j**) and convex hull perimeters (**k**) of CIMs after 1 day in low-adhesion U-bottom well plates in the presence of 20 ng mL^-1^ interferon-α (IFNα), 100 ng mL^-1^ lipopolysaccharide (LPS), 20 ng mL^-1^ IFNγ or 20 ng mL^-1^ IFNγ + 100 ng mL^-1^ LPS. Mean ± s.d., *n* = 7-9 wells per condition, ordinary one-way ANOVA and Tukey’s multiple comparisons test. Scale bar 200 μm. l. Pearson correlation between the convex hull perimeters of CIMs in **j** and BMDMs from Fig. 2a.

**Figure S5.**
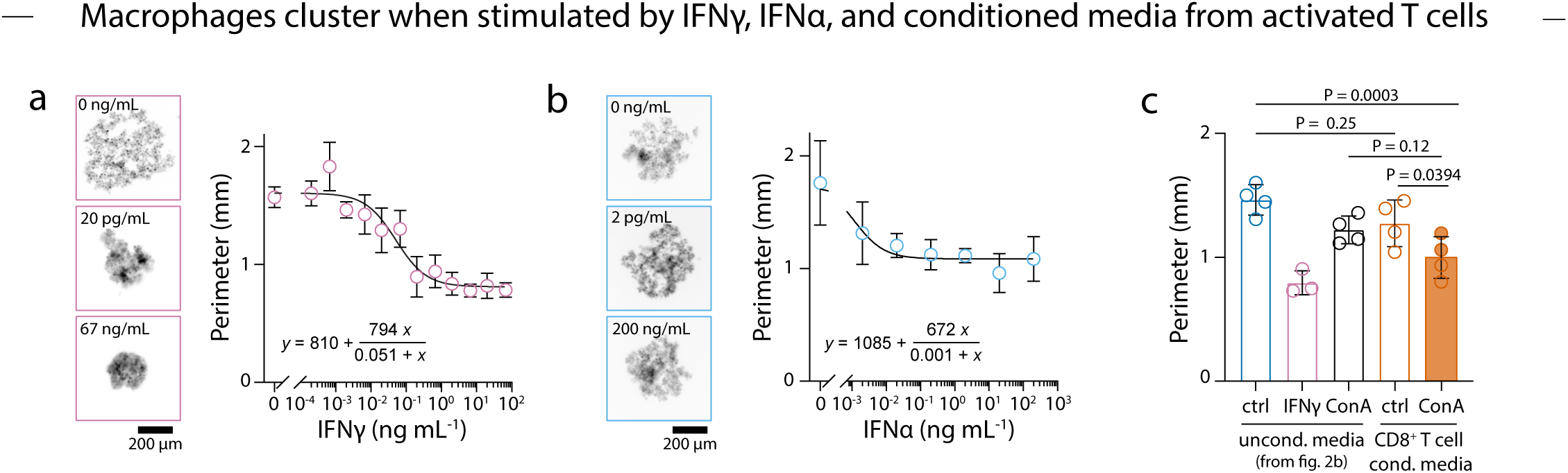
Dose-dependence of IFNγ and IFNα-mediated BMDM clustering and clustering mediated by conditioned medium from activated CD8^+^ T cells a,b. Representative fluorescence images and convex hull perimeters of BMDMs after 1 day on low-adhesion U-bottom well plates with varying concentrations of IFNγ (**a**) or IFNα (**b**). Mean ± s.d., *n* = 5-10 wells per concentration. Scale bar 200 μm. c. Convex hull perimeters of BMDMs after 1 day in low-adhesion U-bottom well plates in the presence of conditioned media from concanavalin-A (ConA)-activated or unactivated mouse CD8^+^ T cells, or in unconditioned media with ConA or IFNγ. Mean ± s.d., *n* = 3 or 4 wells per condition, ordinary one-way ANOVA with Sidak’s multiple comparisons test between indicated groups.

**Figure S6.**
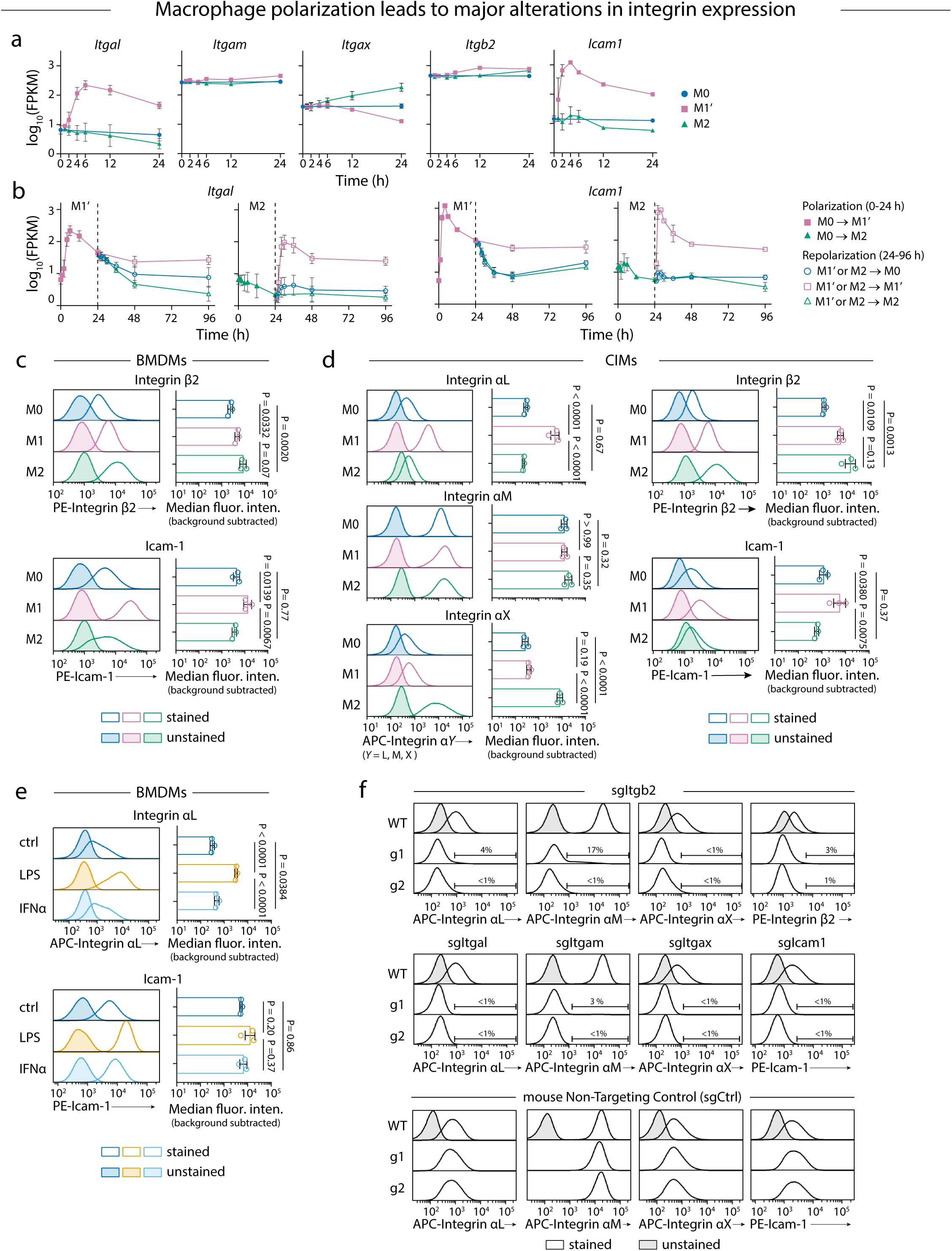
Integrin and Icam-1 expression in polarized macrophages and verification of knockout in gene-edited CIMs a. Kinetic profiles of *Itgal*, *Itgam*, *Itgax*, *Itgb2*, and *Icam1* gene expression in BMDMs cultured in M0, M1’, or M2 medium and harvested at the indicated time points for RNAseq. FPKM = fragments per kilobase million. Mean ± s.d., *n* = 3 biological replicates. b. Kinetic profiles of *Itgal* and *Icam1* gene expression in BMDMs cultured in M1’ or M2 medium for 24 h prior, repolarized in M0, M1’, or M2 medium, and harvested at the indicated time points for RNAseq. Mean ± s.d., *n* = 3 biological replicates. Data for **a** and **b** are from GSE158094 in ref. (37). c-e. Flow cytometry staining of BMDMs cultured for 2 days in M0, M1, or M2 medium and stained with antibodies against integrin β2 and Icam-1 (**c**), differentiated CIMs cultured for 2 days in M0, M1, or M2 medium and stained with antibodies against integrin αL, αM, αX, or β2, or Icam-1 (**d**), and BMDMs cultured for 1 day in medium containing LPS or IFNα and stained with antibodies against integrin αL or Icam-1 (**e**). Histograms show staining of a representative sample and bar graphs report the MFI of stained cells corrected by subtracting the MFI of the unstained culture. Cells were cultured on plastic Petri dishes. Mean ± s.d., *n* = 3 BMDM cultures from different mice (**c,e**) or 3 unique CIM cultures (**d**), ordinary one-way ANOVA and Tukey’s multiple comparisons test on log-transformed MFI values. f. Flow cytometry staining of macrophages differentiated from wild-type (WT) CIM progenitors or from CIM progenitor lines transduced with the indicated sgRNA construct and selected with puromycin. Differentiated CIMs were stained with antibodies against the integrins αL, αM, αX, or β2 or Icam-1. The percentage of cells with fluorescence exceeding that of the unstained WT cells (shaded histograms) is reported for each of the transduced CIM lines.

**Figure S7.**
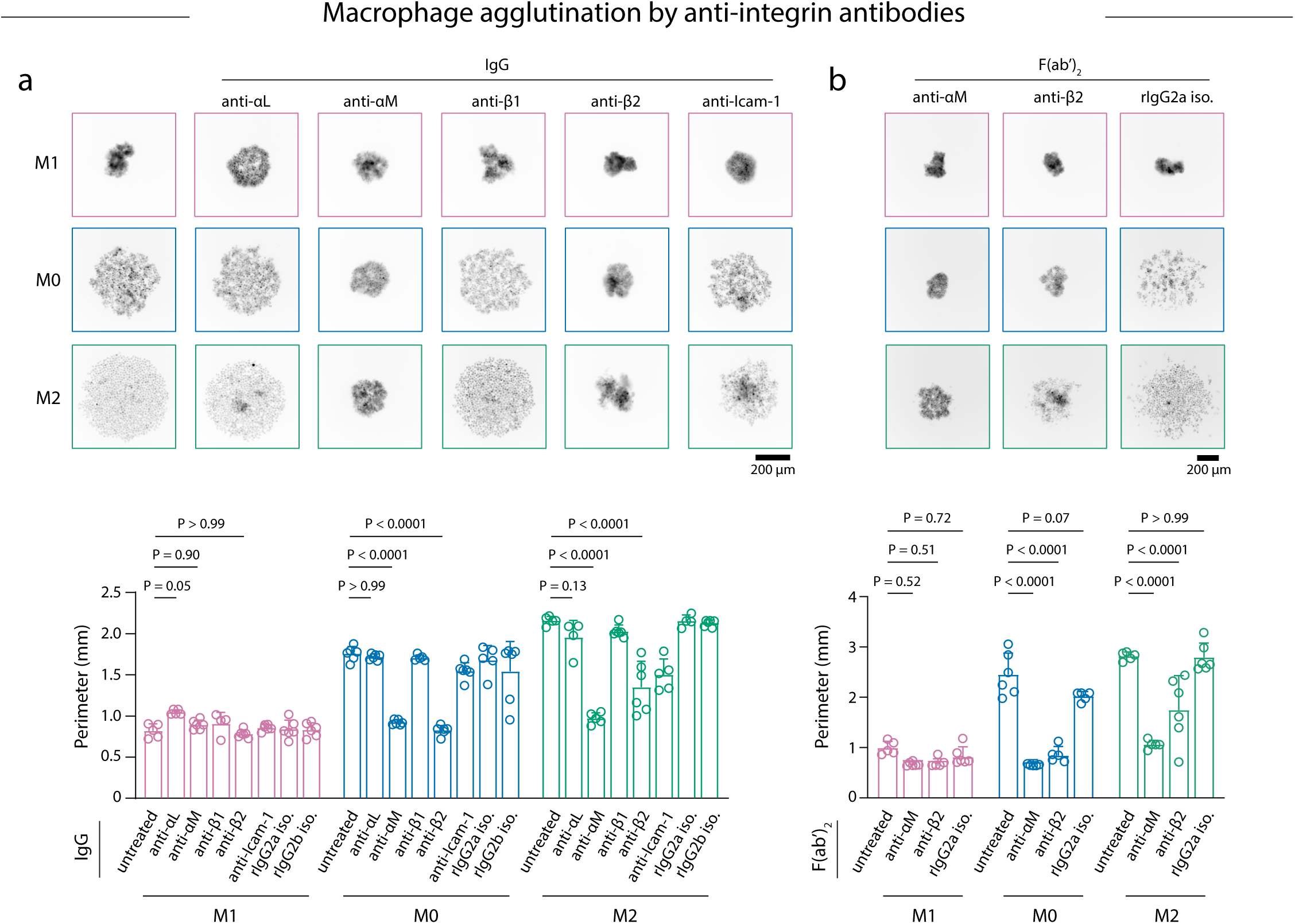
Antibodies and F(ab’)2 fragments targeting abundant macrophage integrins agglutinate BMDMs a,b. Representative images and convex hull perimeters of M0, M1, and M2 BMDMs in the presence of IgG antibodies against integrins αL, αM, β1, or β2, Icam-1, or rat IgG2a or IgG2b isotype control antibodies (**a**) or F(ab’)2 fragments of IgG antibodies against integrin αM or β2 or rat IgG2a isotype control antibody (**b**). Mean + s.d., *n* = 4-6 wells per condition, ordinary two-way ANOVA with Dunnett’s multiple comparisons test between untreated and antibody-treated BMDMs for each polarization condition. Scale bar 200 μm.

**Figure S8.**
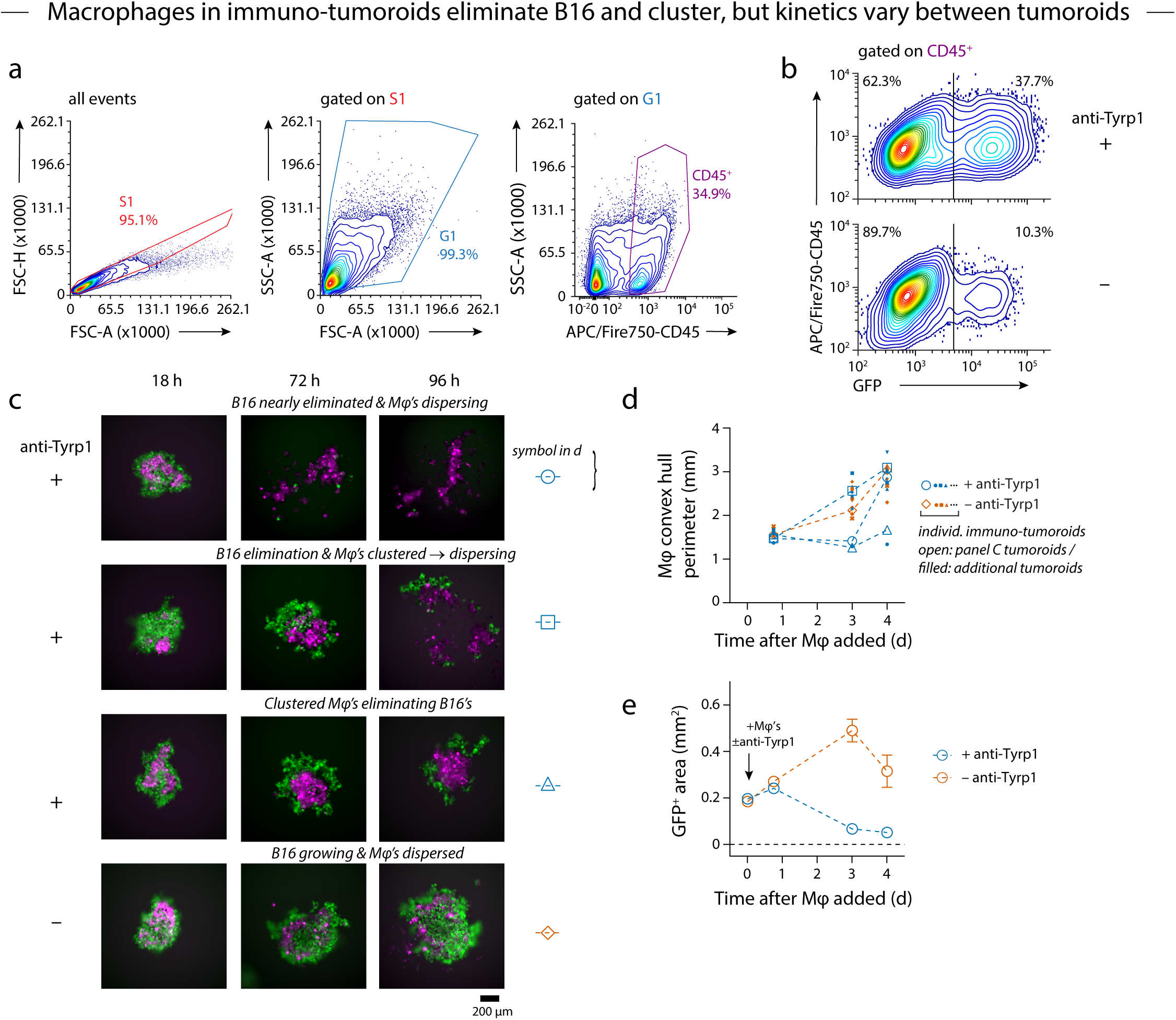
Flow cytometry gating and clustering analysis for macrophages in immuno-tumoroids a,b. Gating strategy for quantifying total macrophages and phagocytic macrophages in flow cytometry. Flow cytometry events were gated on FSC-H vs. FSC-A to discriminate single cells (S1) from doublets, then SSC-A vs. FSC-A to discriminate cells (G1) from debris, and lastly for CD45^+^ macrophages (**a**). CD45^+^GFP^+^ events were considered phagocytic macrophages and CD45^+^GFP^-^ events were considered non-phagocytic macrophages (**b**). c-e. Representative images (**c**) and quantitation of macrophage convex hull perimeters (**d**) and GFP^+^ projected area (**e**) from parallel cultures with fluorescently labeled macrophages performed simultaneously to the flow cytometry analysis in **a**,**b**. For convex hull perimeters, individual tumoroids are shown. For GFP^+^ area, the mean ± standard error of the mean (s.e.m) is shown for *n* = 8 wells per condition. Scale bar 200 μm.

**Figure S9.**
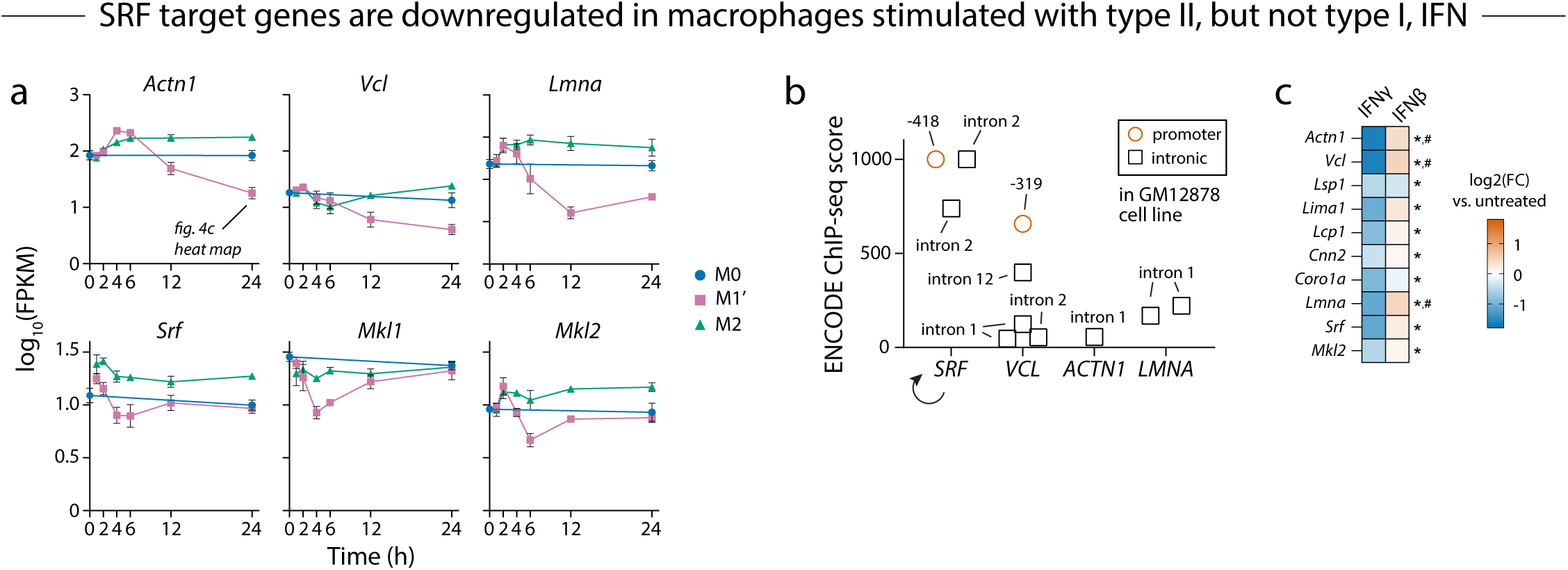
Expression and SRF-regulation of *Vcl*, *Actn1*, and *Lmna* a. Kinetic profiles of *Vcl*, *Actn1*, *Lmna*, *Srf*, *Mkl1*, and *Mkl2* gene expression in BMDMs cultured in M0, M1’, or M2 medium and harvested at the indicated time points for RNAseq. Mean ± s.d., *n* = 3 biological replicates. Data are from GSE158094 ref. (37). b. ChIP-seq score from UCSC Genome Browser/ENCODE for serum-response factor (SRF) binding to promoter or introns of *SRF*, *VCL*, *ACTN1*, and *LMNA* genes in GM12878 lymphoblastoid cells. For promoter peaks, the distance from the translation initiation site is indicated. c. Heat map of fold changes in SRF-related target genes in BMDMs cultured with IFNγ or IFNβ relative to untreated control BMDMs. * and # indicate statistically significant fold change (adjusted P < 0.05) versus untreated for IFNγ and IFNβ, respectively. Data are from GSE 60290.

**Figure S10.**
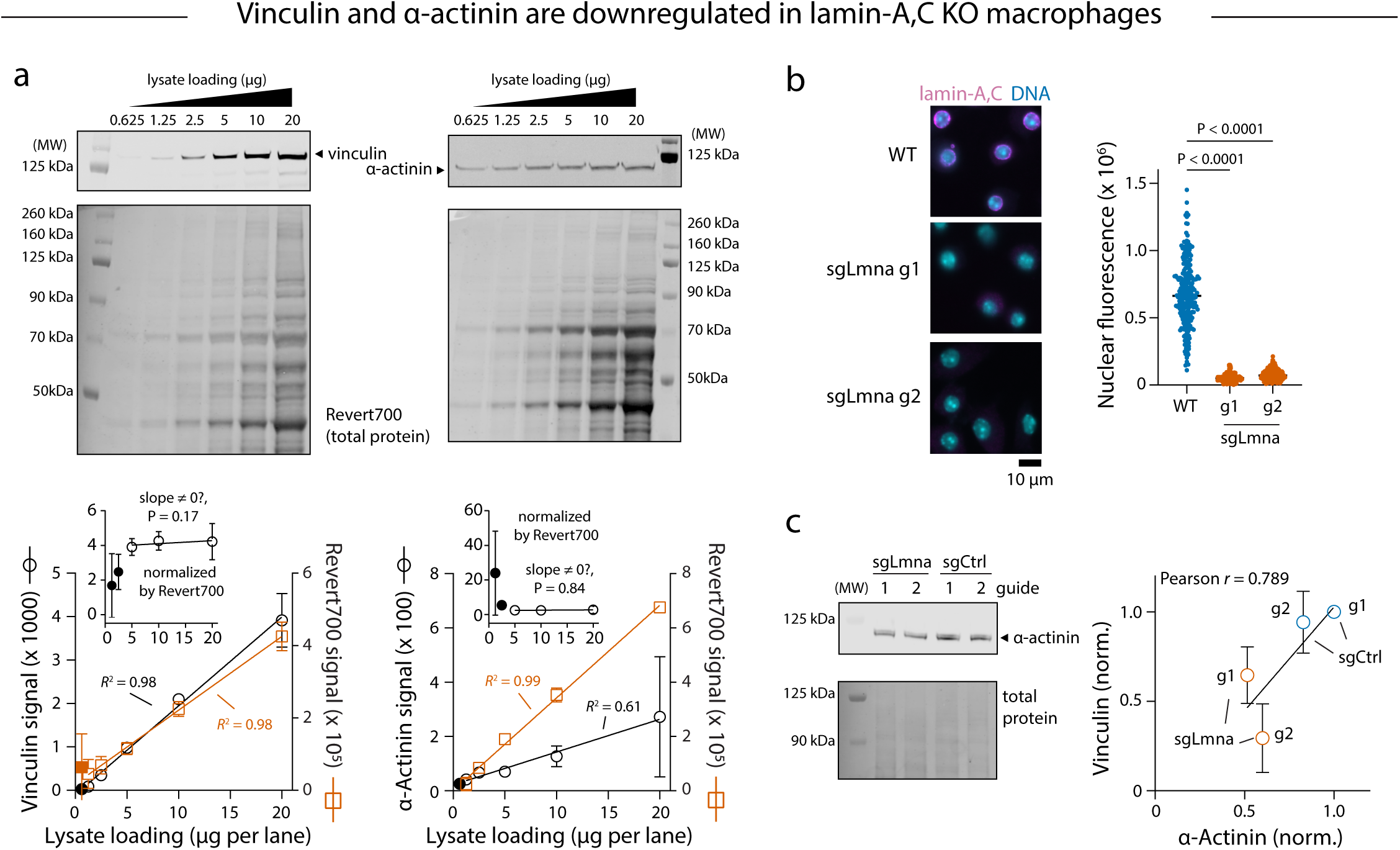
Western blotting of BMDMs and lamin-A,C KO CIMs; immunofluorescence staining of lamin-A,C KO CIMs a. Determination of the linear range of anti-vinculin and anti-α-actinin Western blotting and Revert700 total protein membrane stain by varying lysate loading from 0.625 to 20 μg per lane. Mean ± s.d., lysate amounts were loaded in duplicates. Filled symbols were omitted from regression analyses. The inset plots depict the vinculin or α-actinin band intensity normalized by Revert700 intensity from the same lane and fit to a straight line for lysate concentrations of 5, 10, and 20 μg per lane. Statistical significance was assessed by the extra sum-of-squares F-test with the null hypothesis that the slope equals 0. b. Immunofluorescence staining for lamin-A,C in macrophages differentiated from WT CIM progenitors or progenitors transduced to express sgLmna guides 1 or 2. Scale bar 10 μm. The integrated intensity of lamin-A,C immunofluorescence was calculated for each nucleus. n > 173 cells across 3 fields of view per cell line, outliers were removed by Rout’s method (Q = 1%), Kruskal-Wallis test with Dunn’s multiple comparisons test. c. Immunoblots of lysates from macrophages differentiated from lamin-A,C KO and non-targeting control guide CIMs probed with anti-α-actinin. The α-actinin intensity across the different lysates was normalized by total protein staining to control for loading and then by the intensity of non-targeting control guide 1. The normalized α-actinin intensities are plotted against the normalized vinculin intensity from Fig. 4e.

**Figure S11.**
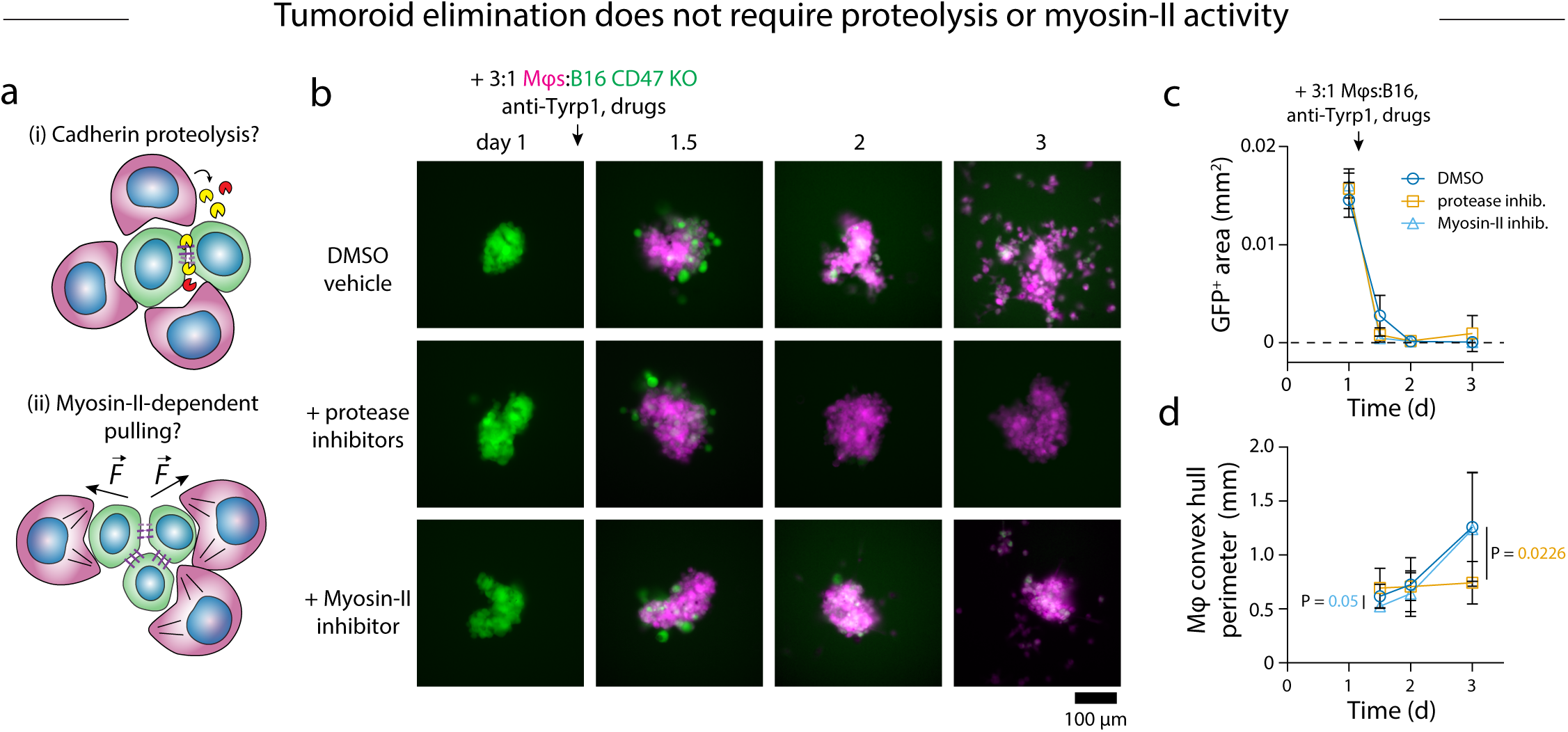
Tumoroid elimination by macrophages does not require proteolysis or myosin-II activity a, Possible mechanisms for disruption of B16 cell-cell adhesions including (i) proteolysis and (ii) myosin-II-dependent pulling forces. b-d. Representative images (**b**), growth curves (**c**), and macrophage convex hull perimeters (**d**) of B16 CD47 KO immuno-tumoroids treated with BMDMs and anti-Tyrp1 added together with DMSO vehicle control, 0.5% v/v protease inhibitor cocktail, or 20 μM NH2-blebb. Mean ± s.d., *n* = 9-10 wells per condition, repeated measures two-way ANOVA with Dunnett’s multiple comparisons test. Scale bar 100 μm.

**Figure S12.**
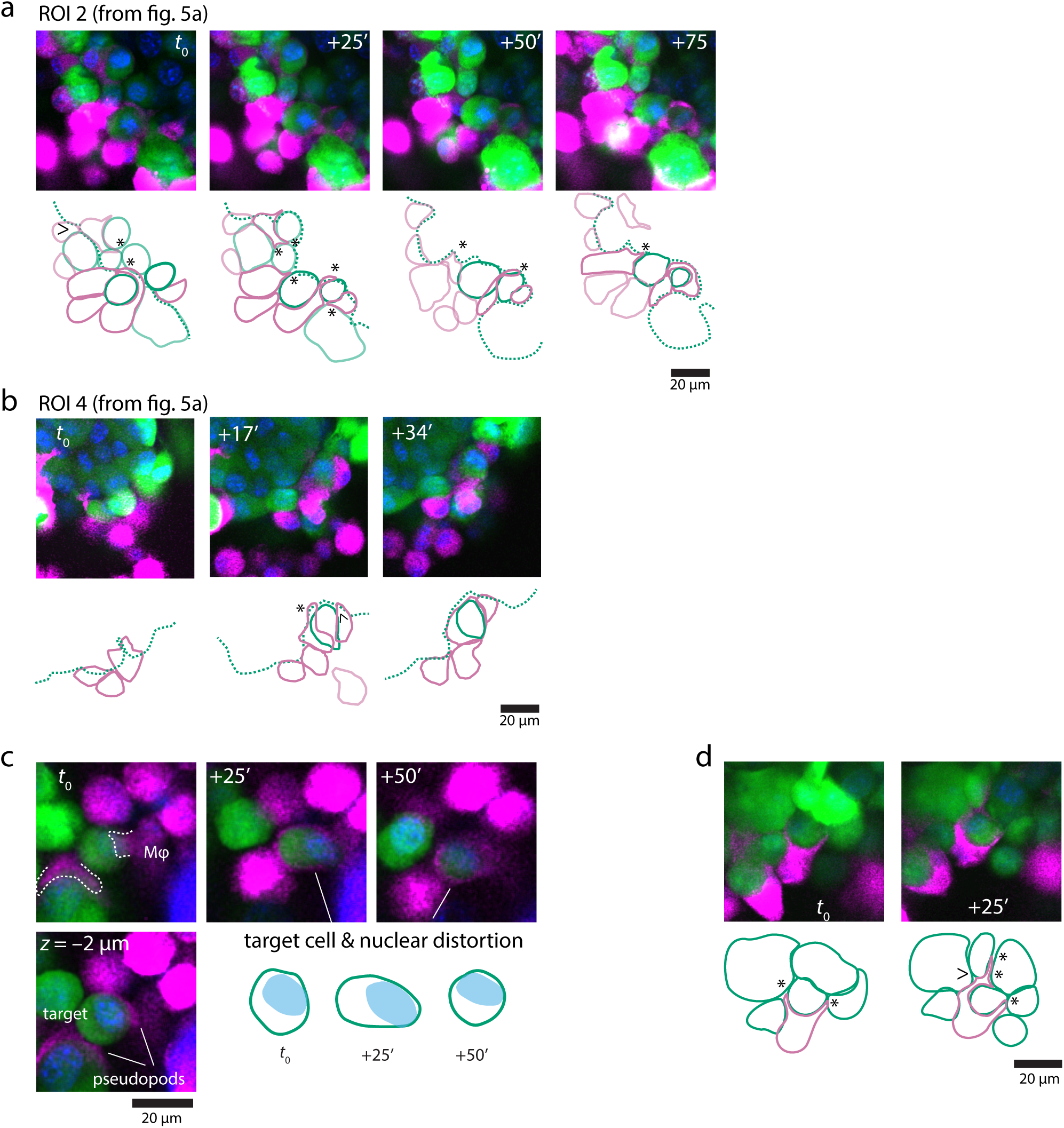
Additional examples of macrophage intrudopodia in B16 spheroids a,b. Zoomed confocal slices of regions of interest 2 (**a**) and 4 (**b**) denoted in **Fig.5a** and accompanying macrophage (magenta) and B16 (green) cell outlines. c. Example phagocytic event in a spheroid with recoverable target cell and nucleus deformation depicted in the cell and nuclear outlines. Dotted white lines show a pseudopod from one macrophage intruding between the target B16 and its neighbor and a second pseudopod spreading over the target B16 that is subsequently engulfed. d. Example of a meandering intrudopod extended by a single macrophage, which did not result in phagocytosis during the experiment. All scale bars 20 μm.

**Figure S13.**
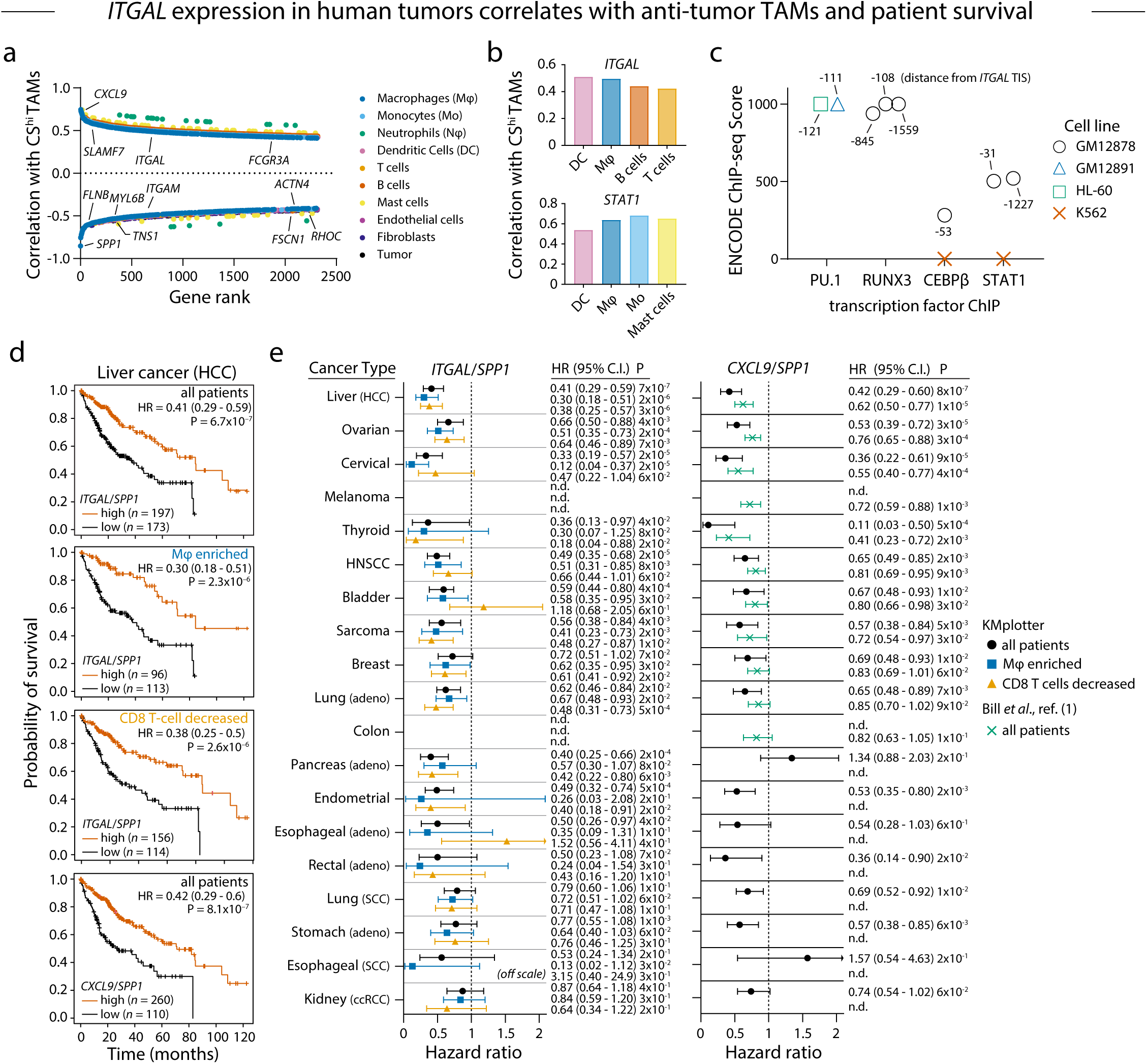
*ITGAL* expression in human tumors correlates with anti-tumor macrophage abundance and patient survival a. Spearman correlation coefficients between gene expression in specific cell types and the abundance of anti-tumor (CS^hi^) macrophages in human head and neck squamous cell carcinoma produced from data reported in ref. (1) . Correlation coefficients are arranged by rank. CS^hi^ TAMs refers to tumor-associated macrophages with a high ratio of *CXCL9* expression to *SPP1* expression. Other genes with macrophage expression levels that correlate with CS^hi^ TAM abundance and that encode proteins implicated in phagocytosis or actin crosslinking and contractility are indicated. b. Correlation coefficients for *ITGAL* and *STAT1* from **a** in multiple cell types. c. ChIP-seq score from UCSC Genome Browser/ENCODE for transcription factors PU.1, CEBPβ, RUNX3, and STAT1 binding to the promoter region of *ITGAL* in the indicated human cell lines. The value accompanying each data point is the from the translation initiation site (TIS). d,e. Kaplan-Meier survival plots for human liver cancer (**d**) as a representative cancer type and summary plots of hazard ratios (HR) across additional types of human cancers (**e**). Patients were stratified based on the ratio of *ITGAL* to *SPP1* expression (top three plots in **d** and left plot in **e**) or the ratio of *CXCL9* to *SPP1* expression (bottom plot in **d** and right plot in **e**). *ITGAL*/*SPP1* ratio analyses were performed separately for all patients, for patients with tumors enriched in macrophages, and for patients with tumors having low numbers of CD8^+^ T cells using options available in the KMplot pan-cancer RNAseq webtool. *CXCL9*/*SPP1* ratio analyses were performed for all patients using KMplot and are shown together with data from ref. (1) to demonstrate comparable effects on survival in both sets of patient data.

## Supplementary Tables 1-3

**Table S1:** Macrophage polarization transcriptomic datasets analyzed for expression of genes encoding adhesion receptors and cytoskeleton.

| **Dataset** | **Polarization** | **Method** | **Reference:** |
| --- | --- | --- | --- |
| GS158094 | M1': 25 ng mL<sup>-1</sup> IFNγ + 100 ng mL<sup>-1</sup> LPS<br>M2: 25 ng mL<sup>-1</sup> IL-4<br>1,2,4,6,12,24 h | RNAseq | (37) |
| GSE69607 | M1': 20 ng mL<sup>-1</sup> IFNγ + 100 ng mL<sup>-1</sup> LPS<br>M2: 20 ng mL<sup>-1</sup> IL-4<br>24 h | Microarray | (76) |
| GSE60290 | M1: 10 U mL<sup>-1</sup> IFNγ<br>IFNβ: concentration not specified<br>18 h | Microarray | (77) |

**Table S2:**
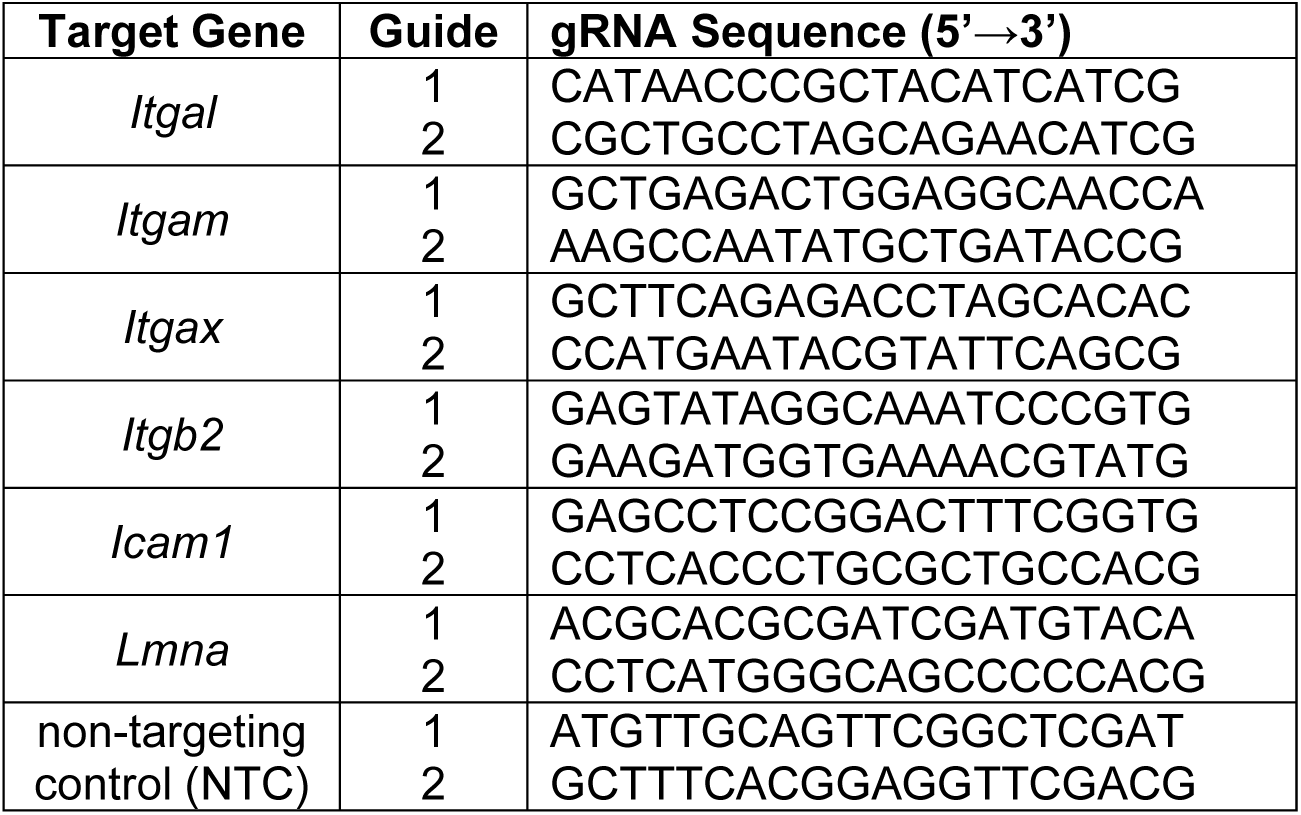
Guide RNA sequences.

**Table S3:** Antibody details.

| Antibody (Isotype) | Fluorophore | Clone | Supplier | Cat. # | Lot #(s) | Conc<br>µg/mL or<br>dilution |
| --- | --- | --- | --- | --- | --- | --- |
| **Flow cytometry characterization of bone marrow-derived macrophages and conditionally immortalized macrophages (Fig.S2a)** |  |  |  |  |  |  |
| Rat anti-mouse CD45 (IgG2b,κ) | APC/Cy7 | 30-F11 | BioLegend | 103116 | B372054 | 2.5 |
| Rat anti-mouse CD11b (IgG2b,κ) | APC | M1/70 | BioLegend | 101212 | B384396 | 2.5 |
| Rat anti-mouse CD115 (IgG2a,κ) | PE/Cy7 | AFS98 | BioLegend | 135523 | B347333 | 1.25 |
| Rat anti-mouse F4/80 (IgG2a,κ) | PE | BM8 | BioLegend | 123109 | B262795 | 10 |
| Rat anti-mouse Ly6C (IgG2c,κ) | PE | HK1.4 | BioLegend | 128007 | B239361 | 2.5 |
| Rat anti-mouse CD172a (IgG1,κ) | PE/Cy7 | P84 | BioLegend | 144007 | B359167 | 2.5 |
| Rat anti-mouse CD64 (IgG2a,κ) | PE | S180117D | BioLegend | 161003 | B363525 | 2.5 |
| **M1 and M2 marker flow cytometry (Fig.S2b,c)** |  |  |  |  |  |  |
| Rat anti-mouse I-A/I-E (MHCII) (IgG2b,κ) | APC | M5/114.15.2 | BioLegend | 107613 | B322689 | 2.5 |
| Rat anti-mouse CD206 (MMR) (IgG2a,κ) | PE/Cy7 | C068C2 | BioLegend | 141719 | B253128 | 2.5 |
| **Integrin expression profiling flow cytometry (Fig.2d, Fig.S6c-f)** |  |  |  |  |  |  |
| Rat anti-mouse CD11a (integrin αL) (IgG2a,κ) | APC | M17/4 | BioLegend | 101119 | B266513 | 10 |
| Rat anti-mouse/human CD11b (integrin αM) (IgG2b,κ) | APC | M1/70 | BioLegend | 101211 | B312599 | 5 |
| Armenian hamster anti-mouse CD11c (integrin αX (IgG) | APC | N418 | BioLegend | 117309 | B339314 | 5 |
| Rat anti-mouse CD18 (integrin β2) (IgG2a,κ) | PE | M18/2 | BioLegend | 101407 | B373658 | 20 |
| Rat anti-mouse CD54 (Icam-1) (IgG2b,κ) | PE | YN1/1.7.4 | BioLegend | 116107 | B314625 | 0.6 |
| **Tumoroid macrophage surface marker profiling flow cytometry (Fig.3b-c, Fig.S8a,b)** |  |  |  |  |  |  |
| Rat anti-mouse CD45 (IgG2b,κ) | APC/Fire750 | 30-F11 | BioLegend | 103153 | B418348 | 2.5 |
| Rat anti-mouse I-A/I-E (MHCII) (IgG2b,κ) | APC | M5/114.15.2 | BioLegend | 107613 | B374585 | 2.5 |
| Rat anti-mouse CD11a (integrin αL) (IgG2a,κ) | PE | M17/4 | BioLegend | 101107 | B382505 | 5 |
| Rat anti-mouse CD54 (Icam-1) (IgG2b,$\kappa$) | BV421 | YN1/1.7.4 | BioLegend | 116141 | B425621 | 0.6 |
| **Western Blotting (WB) and Immunofluorescence (IF) Microscopy (Fig.4d,e, Fig.S10)** |  |  |  |  |  |  |
| Rabbit anti-α-actinin (IgG) | - | D6F6 | CST | 6487 | 4 | 1:1000 |
| Rabbit anti-vinculin (IgG) | - | E1E9V | CST | 13901 | 9 | 1:1000 |
| Mouse anti-lamin-A,C (IgG2a,$\kappa$) | - | 4C11 | CST | 4777 | 4,5 | 1:1000<sup>WB</sup><br>1:100<sup>IF</sup> |
| Goat anti-mouse IgG [H+L] | IRDye800CW | Polyclonal | LiCor | 925-32210 | D30228-01 | 1:15,000 |
| Donkey anti-rabbit IgG [H+L] | IRDye800CW | Polyclonal | LiCor | 925-32213 | D30627-11 | 1:15,000 |
| Donkey anti-mouse IgG [H+L] | Alexa Fluor Plus 647 | Polyclonal | Invitrogen | A32787 | YB363609 | 10 |
| Donkey anti-rabbit IgG [H+L] | Alexa Fluor Plus 647 | Polyclonal | Invitrogen | A32795 | XJ359900 | 2 |
| **Blockade (all functional grade, unconjugated, low endotoxin, azide-free) (Fig.S7)** |  |  |  |  |  |  |
| Rat anti-mouse CD11a (integrin αL) (IgG2a,$\kappa$) | - | M17/4 | BioLegend | 101118 | B381281 | 20 |
| Rat anti-mouse/human CD11b (integrin αM) (IgG2b,$\kappa$) | - | M1/70 | BioLegend | 101248 | B346553 | 20 |
| Rat anti-mouse (integrin β1) (IgG2a,$\kappa$) | - | KMI6 | BioXCell | BE0232 | 787622F1 | 20 |
| Rat anti-mouse CD18 (integrin β2) (IgG2a,$\kappa$) | - | M18/2 | BioLegend | 101418 | B398788 | 20 |
| Rat anti-mouse CD54 (Icam-1) (IgG2b,$\kappa$) | - | YN1/1.7.4 | BioLegend | 116123 | B371691 | 20 |
| Rat IgG2a,$\kappa$ isotype control | - | RTK2758 | BioLegend | 400544 | B340640 | 20 |
| **Opsonization of Tumors, Tumoroids, Spheroids, B16 suspensions, and polystyrene beads (Fig.1a-d, Fig.S1, Fig.S3, Fig.4g-h, Fig.5a,b Fig.S11, Fig.S12)** |  |  |  |  |  |  |
| Mouse anti-human/mouse Tyrp1 (IgG2a,$\kappa$) | - | TA99 | BioXCell | BE0151 | 715421A1 | 20 |
| Rabbit anti-streptavidin | - | Polyclonal | Sigma | S6930 | 097M4817 V | 1:400 |

## Notes

### Competing Interest Statement

The authors have declared no competing interest.

